# Memory B cell repertoire for recognition of evolving SARS-CoV-2 spike

**DOI:** 10.1101/2021.03.10.434840

**Authors:** Pei Tong, Avneesh Gautam, Ian Windsor, Meghan Travers, Yuezhou Chen, Nicholas Garcia, Noah B. Whiteman, Lindsay G.A. McKay, Felipe J.N. Lelis, Shaghayegh Habibi, Yongfei Cai, Linda J. Rennick, W. Paul Duprex, Kevin R. McCarthy, Christy L. Lavine, Teng Zuo, Junrui Lin, Adam Zuiani, Jared Feldman, Elizabeth A. MacDonald, Blake M. Hauser, Anthony Griffths, Michael S. Seaman, Aaron G. Schmidt, Bing Chen, Donna Neuberg, Goran Bajic, Stephen C. Harrison, Duane R. Wesemann

## Abstract

Memory B cell reserves can generate protective antibodies against repeated SARS-CoV-2 infections, but with an unknown reach from original infection to antigenically drifted variants. We charted memory B cell receptor-encoded monoclonal antibodies (mAbs) from 19 COVID-19 convalescent subjects against SARS-CoV-2 spike (S) and found 7 major mAb competition groups against epitopes recurrently targeted across individuals. Inclusion of published and newly determined structures of mAb-S complexes identified corresponding epitopic regions. Group assignment correlated with cross-CoV-reactivity breadth, neutralization potency, and convergent antibody signatures. mAbs that competed for binding the original S isolate bound differentially to S variants, suggesting the protective importance of otherwise-redundant recognition. The results furnish a global atlas of the S-specific memory B cell repertoire and illustrate properties conferring robustness against emerging SARS-CoV-2 variants.

## INTRODUCTION

Coronavirus (CoV) disease 2019 (COVID-19) caused by the severe acute respiratory syndrome (SARS) CoV-2 virus has rapidly become a pandemic of historic effect. Although vaccines have been developed in record time, new variants continue to emerge and threaten to evade immune responses. We need to understand immune recognition of SARS-CoV-2, especially as stored in B cell memory, to illuminate the requirements for broad protective immunity in humans. We focus on B cells, because antibodies, a key part of the immune defense against most viruses, are sufficient to protect against SARS-CoV-2 infection in animal models (*1, 2*).

Antibodies are both soluble effector molecules and the antigen-receptor component of the B cell receptor (BCR). BCRs evolve enhanced pathogen binding through immunoglobulin (Ig) gene somatic hypermutation (SHM) and selection in lymphoid tissue germinal centers (GCs), leading to antibody affinity maturation (*3*) and generation of both antibody-secreting plasma cells (PCs) and memory B cells. Higher avidity interactions encourage terminal differentiation of B cells into PCs; memory B cells frequently have lower avidity but more cross-reactive specificities (*4*).

Both PC-derived secreted antibody and memory B cells supply immune memory to prevent repeat infection, but with non-redundant roles. Secreted antibodies can prophylactically thwart pathogen invasion with fixed recognition capability, while memory B cells harbor expanded pathogen recognition capacity and can differentiate quickly into PCs to contribute dynamically to the secreted antibody repertoire (*4*). Moreover, memory B cells retain plasticity to adapt to viral variants through GC re-entry and SHM-mediated evolution (*5*).

The viral spike (S) glycoprotein binds ACE2 on host cells and mediates viral fusion with the host (*6*). Its fusogenic activity depends on a furin-mediated cleavage, resulting in N-terminal S1 and C terminal S2 fragments (*7*) and on a subsequent cleavage of S2 mediated either by cathepsins or by a serine protease, TMPRSS2 (*8*). The S glycoprotein is the principal neutralizing antibody target and the focus of most vaccines. SARS-CoV-2 S antibodies decline with time (*9, 10*) and can lose reactivity to emerging variants (*11*). Antibodies cloned from memory B cells target the S glycoprotein in redundant as well as unique ways, indicating cooperative and competitive recognition (*12–17*). Many of these antibodies have been identified and characterized; their positions within the distribution of practical cooperative recognition of SARS-CoV-2 S within the human memory B cell repertoire have not. Moreover, the recognition reach of memory B cells induced by one SARS-CoV-2 strain toward evolving stains across the major epitopic regions has not yet been defined.

We present here an unbiased global assessment of the distribution of memory B-cell encoded antibodies among cooperative and competitive recognition clusters on the SARS-CoV-2 S glycoprotein and assess features that direct their collaborative robustness against emerging SARS-CoV-2 variants. In a comprehensive competition analysis of 152 monoclonal antibodies (mAbs) from 19 subjects for binding with trimeric S ectodomain, we have identified 7 recurrently targeted competition groups -- three for antibodies with epitopes on the receptor-binding domain (RBD), two for epitopes on the N-terminal domain (NTD), and two for S2 epitopes. We show that these groups represent the major practical antibody footprints, with rare antibodies outside them. We map the clusters onto the S glycoprotein by including previously characterized antibodies and new cryo-EM determined structures. Ig repertoire analysis indicates both divergent and convergent clones with the competition groups.

Antibodies mapped to RBD-2 and NTD-1 were the most potent neutralizers, while the S2-1 group has the greatest recognition breadth across CoVs. The emerging SARS-CoV-2 variants, particularly the South Africa strain, strongly affected the antibodies in one of the RBD and one of the NTD clusters. The mutations in those variants differently influenced affinity of antibodies within a competition group, indicating that the depth of otherwise redundant mAbs to a given S variant confers recognition breadth for dynamically mutating S.

## RESULTS

### Monoclonal antibody (mAb) isolation

To identify the general pattern of SARS-CoV-2 S recognition by memory B cells in convalescent subjects, we sorted single CD19^+^ CD27^+^ IgG^+^ B cells recognizing soluble prefusion-stabilized S trimer (Fig. 1A, Fig. S1) from 19 individuals with a history of COVID-19 (Data S1). Because less is known about S-reactive antibodies that bind outside the RBD region, we also sorted S-reactive B cells that did not bind RBD from 3 individuals. S-reactive B cells made up 0.2% (0.07%-0.4%) of the total B cell population (Fig. 1A left panel), with RBD-binding cells representing about a quarter of S-reactive IgG^+^ B cells (Fig. 1A right panel) consistent with prior work (*18*).

**Fig. 1.**
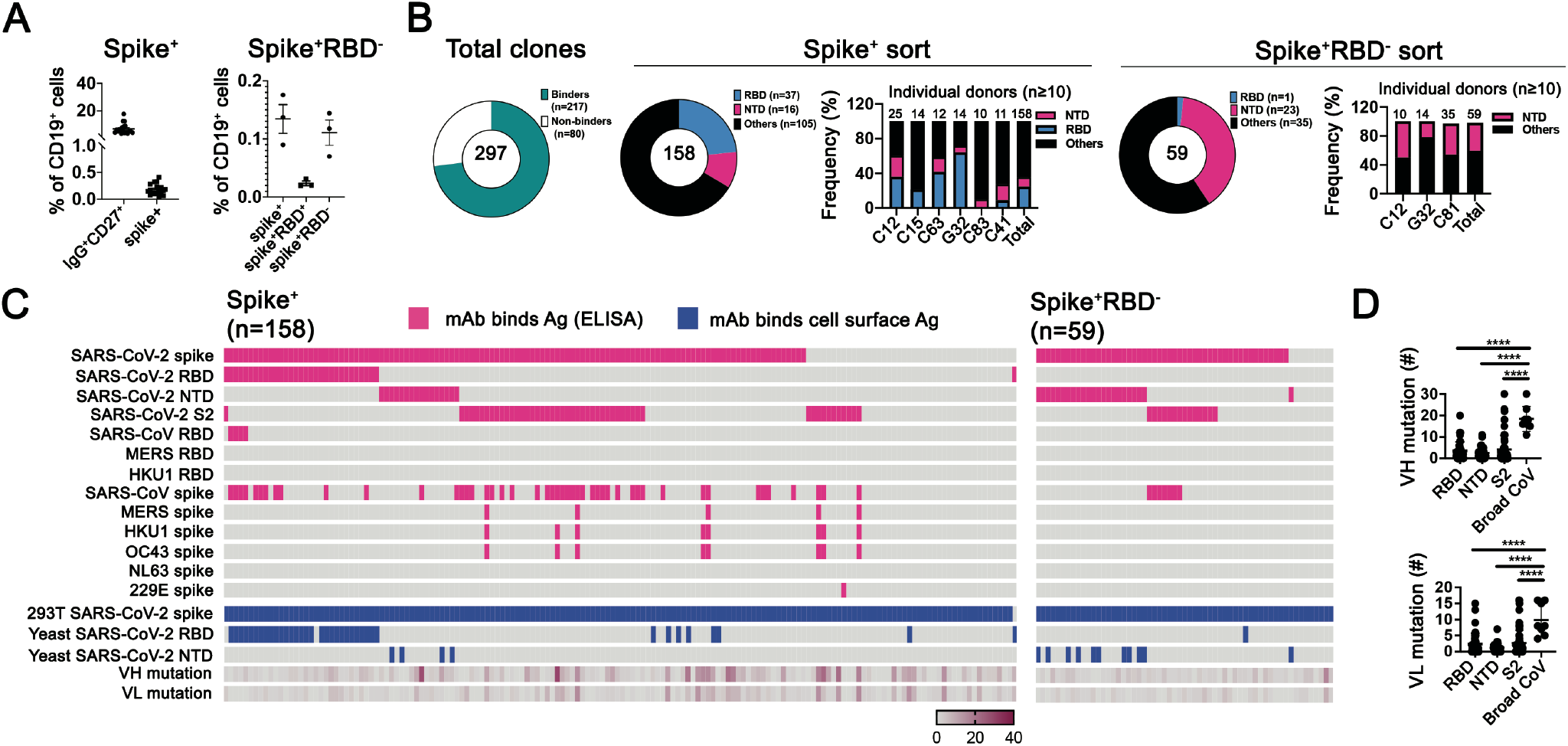
SARS-CoV-2 surface glycoprotein (spike) specificities of memory B cells from convalescent subjects. **(A)** Cells recovered from two sorting strategies, shown in dot plots as percentages of total CD19^+^ cells. Left: IgG^+^CD27^+^ cells from 18 donors (one dot per donor) and the subset of those that expressed spike-binding BCRs. Right: cells from 3 donors expressing spikebinding BCRs and sorted to recover principally those that did not bind recombinant receptor-binding domain (RBD). Sorting protocols as described in Methods and shown in Fig. S1. **(B)** Summary of all antibodies (expressed as recombinant IgG1) screened by ELISA (with recombinant spike ectodomain trimer) and cell-surface expression assays (both 293T and yeast cells). Total numbers in the center of each of pie chart; numbers and color codes for the indicated populations shown to next to each chart. To the right of the charts for the two alternative sorting strategies are bar graphs showing frequencies of SARS-CoV-2 RBD and NTD binding antibodies for those subjects from whom at least 10 paired-chain BCR sequences were recovered. **(C)** Binding to a panel of spike proteins and SARS-CoV-2 subdomains, listed on the left, as determined by both ELISA (with recombinant spike ectodomain) and by association with spike expressed on the surface of 293T cells or with RBD or NTD expressed on the surface of yeast cells, for cells sorted just for spike binding (left) and for those sorted for positive spike binding but no RBD binding (right). The rows with pink highlighting are from the ELISA screen; those with blue highlighting, from the cell-based screens. Each short section of a row represents an antibody. The rows labeled VH mutation and VL mutation are heat maps of counts (excluding CDR3) from alignment by IgBLAST, with the scale indicated. **(D)** Dot plots of heavy-and light-chain somatic mutation counts in antibodies that bound RBD, NTD, S2, and a “broad CoV group” that included MERS, HKU1, and OC43. The significantly higher numbers of mutations in the last group suggest recalled, affinity matured memory from previous exposures to seasonal coronaviruses. ****P < 0.0001; one-way ANOVA followed by Tukey’s multiple comparison. Horizontal lines show mean ± SEM.

### mAb binding

We cloned cDNAs encoding Ig heavy (H) and light (L) chains from individual, sorted memory B cells into human IgG1 and kappa or lambda vectors and expressed them in HEK 293T cells. We detected IgG in 255 of the culture supernatants, which we used to screen for binding of SARS-CoV-2 S (Fig. 1 and fig. S2). Of the 255 IgGs, 216 bound SARS-CoV-2 S expressed by HEK 293T cells, as assayed by flow cytometry (157 from the S^+^ sorting, 59 from S^+^/RBD^-^ sorting) (fig. S2A) and 166 of the 216 bound recombinant SARS-CoV-2 S, as assayed by ELISA (116 from the S^+^ sorting, 50 from the S^+^RBD^-^ sorting).

We estimated, by ELISA and, where possible, yeast display of the subdomains (fig. S2B and C), the proportion of mAbs that bound to RBD, NTD, and S2. Recombinant and yeast-displayed S2 protein could have any of several conformations, and all or parts of the polypeptide chain might be disordered; antibodies that bound S2 on ELISA plates might therefore tend to recognize linear epitopes or even the S2 post-fusion conformation. Indeed, most of the S2-binding antibodies had relatively low ELISA-determined affinities for intact, recombinant, prefusion S, although a few bound more tightly to S expression on the surface of 293T cells (fig. S2D). Of the 157 S-reactive mAbs sorted with SARS-CoV-2 stabilized S trimer, a total of 37 (23%) were RBD-specific as assayed by ELISA, by yeast display, or both (Fig. 1B). We detected 16 (10%) mAbs that bound the NTD and 49 (31%) that bound recombinant S2 (Fig. 1C). Eleven of the 49 S2 binders bound cell-surface-expressed, but not ELISA-based SARS-CoV-2 S.

We also assessed mAbs by ELISA for cross-reactivity to other CoV S glycoproteins. Those of SARS (GenBank: MN985325.1), MERS (GenBank: JX869059.2), and common cold β-CoVs HKU1 (GenBank:Q0ZME7.1) and OC43 (GenBank: AAT84362.1) have sequences with 75.8%, 28.6%, 25.1%, and 25.5 % amino-acid identity, respectively, with SARS-CoV-2 S; the more distantly related common cold a-CoVs, NL63 (AAS58177.1) and 229E (GenBank: AAK32191.1), just 18.3% and 20.2 %. Of the 157 S ectodomain-sorted mAbs, 47 (29.9%) bound to SARS-CoV S and 8 to other β-CoV S glycoproteins. These 8 cross-reactive antibodies have higher mutation levels than do RBD, NTD, and the other S2-binding mAbs from our cohort (Fig. 1C and D). Among the 59 S-binding mAbs cloned from the S^+^RBD^-^ sorted memory B cells, ELISA detected 23 (39%) mAbs that bound NTD (11 of which also bound NTD on yeast), 14 (23.7%) that bound S2, of which 7 (11.9%) cross-reacted with SARS-CoV S. One mAb bound RBD (Fig. 1B and C).

### Global competition analysis defines seven epitopic regions on S

We used a competition ELISA to determine pairwise overlaps of the 105 antibodies in our panel for which we could detect signal at 1 μg/mL. By adding a biotinylated version of each mAb (1 μg/mL) together with 100-fold excess of each of the other mAbs individually into ELISA plates pre-coated with pre-fusion-stabilized SARS-CoV-2 S (*19*), we could detect competition of mAbs with up to 100-fold differences in affinity. We also included 15 published mAbs with known structures as references (fig. S3A).

We identified seven major clusters of competing mAbs—three RBD clusters, two NTD clusters, and two S2 clusters (fig. S3A). The three RBD clusters overlapped to varying extents, as expected for sites on a relatively small domain. Asymmetric competition (one mAb blocks binding of another, but the second does not block the first) occurred when one had much higher affinity than the other -- e.g., S309 (*20*), which binds more tightly than do most of the RBD-1 mAbs we isolated. The clusters define relatively broad epitopic regions, as the footprints of two antibodies within a cluster might not overlap with each other but both might overlap with the footprint of a third (e.g., REGN10933, REGN10987, both of which competed with CC12.1, although they have completely distinct footprints at either end of the RBD receptor binding motif, (RBM) (*21*). Some crosstalk between clusters is also evident (e.g. C93D9, which bound the RBD, blocked both RBD-2 and NTD-1 mAbs). The published 4-8, 4A8 and COVA1-22 mAbs (*12, 13, 22*), which have been shown to bind the NTD, compete with each other and map to NTD-1. NTD-2 mAbs cluster distinctly from NTD-1, indicating minimal spatial overlap of these two NTD regions. One NTD-1 mAb (C81H11) competed strongly with antibodies from S2-1, and a second could be assigned on the basis of competition either to NTD-1 or to S2-1, suggesting structural adjacency of at least some sites in these two clusters (fig. S3A). Several segments of S2 are in contact with either RBD or NTD, some with differential exposure depending on whether the RBD is “up” or “down”.

Thirty-six mAbs in the ELISA competition analysis cross-reacted with SARS-CoV. These mapped mostly to the RBD-1 (11 mAbs) and S2-1 (17 mAbs) clusters. Four mAbs that mapped to S2-1 (C15C3, C7A4, C7A9 and G32Q1) also bound the common cold β-CoVs, and two of these (C7A9, C15C3) also bound MERS S. Thus, S2-1 mAbs appear to recognize a region of S2 conserved among SARS-CoV-2, SARS-CoV, MERS, HKU1 and OC43, as shown previously for S2 antibodies (*23, 24*).

Isolation of a single mAb (C12B3) that bound S2 but did not map to any of the seven major clusters suggests that the immune system may target additional regions of S2, but that those responses are subdominant.

### Memory B cells dominant across individuals in natural infection

We probed the relative distribution of epitopes recognized by SARS-CoV-2 specific memory B cells in the population represented by our cohort by ELISA-based and cell surface-based assays. We clustered all the S^+^ mAbs (Fig. 2) and S^+^RBD^-^ mAbs from a separate sorting step (fig. S3B and C). In the former set, comprising 73 mAbs that bound strongly enough for the ELISA competition assay, the order of epitopic region frequencies was RBD-1 (27.4%), S2-1 (19.2%), NTD-1 (17.8%), RBD-2 (15.1%), NTD-2 (8.2%) (Fig. 2A). There were 36 more mAbs that had insufficient affinity for ELISA competition but that bound cell surface SARS-CoV-2 S (Fig. 2B and fig. S4A). To map ELISA-insufficient mAbs to the 7 clusters, we mixed the biotinylated antibodies with blocking antibodies selected from the ELISA-mapped competition assays, incubated the mixture with cells expressing SARS-CoV-2 S, and recorded the mean value florescent intensity (MFI) to calculate the blocking strength. We used twenty mAbs (2 RBD-3, 4 RBD-2, 4 RBD-1, 2 NTD-2, 4 NTD-1, 2 S2-1 and 2 S2-2) from the ELISA-mapped competition clusters as blocking antibodies, a non-COVID-19 related blocking antibody as negative control, and self-blocking as positive control (Fig. 2B and S3C). We used an additional 118 mAbs (distributed across the 7 clusters) to map the 13 mAbs that failed to compete with the initial 20 (fig. S4B).

**Fig. 2.**
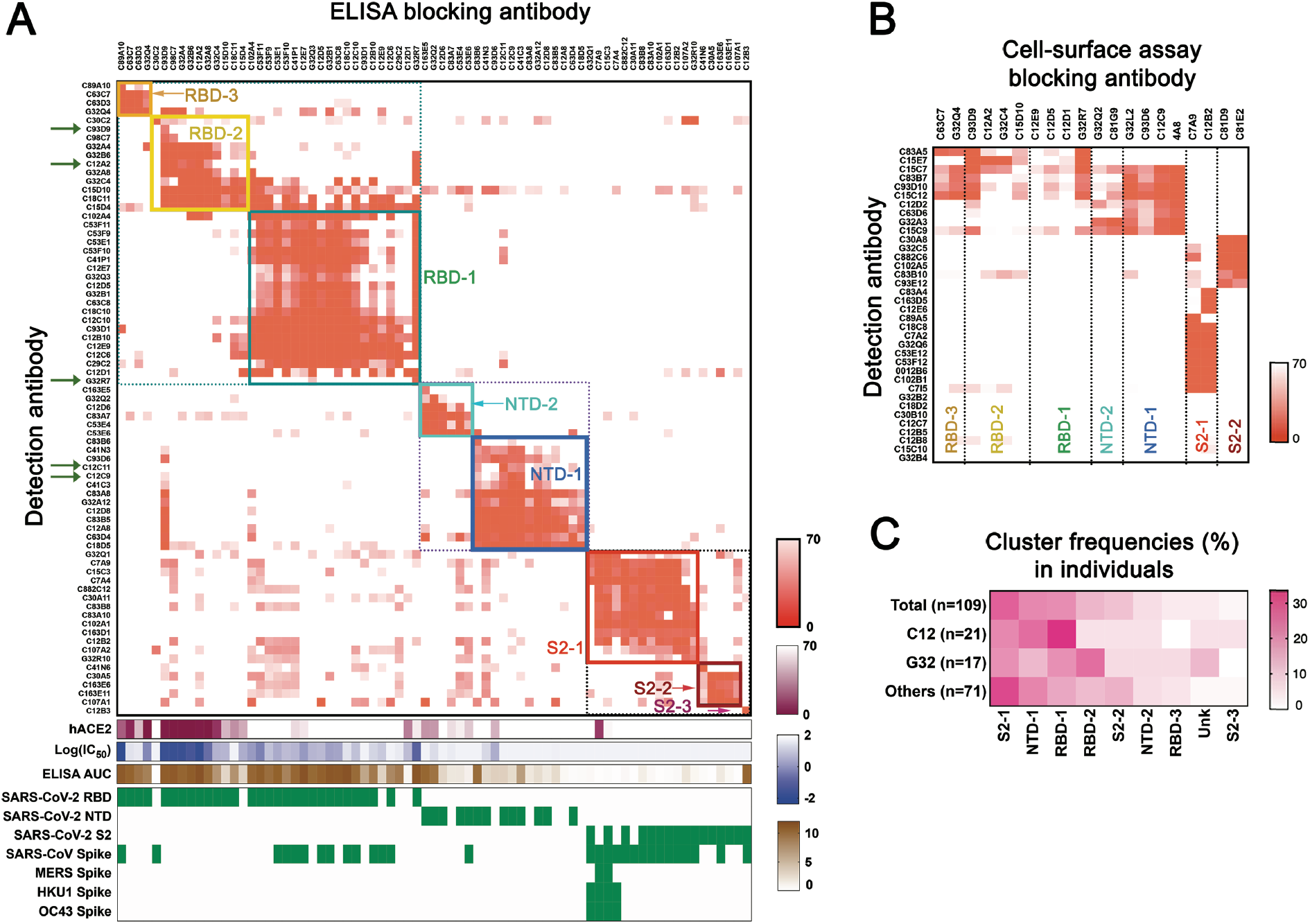
Competition epitope mapping. (**A**) Cross competition matrix for 73 antibodies from the spike^+^ sort in Fig. 1 with affinity sufficient for detection by ELISA. Blocking antibodies (columns) added at 100 μg/ml; detection antibodies (rows), at 1 μg/ml. Intensity of color shows strength of blocking, from 0 signal (complete blocking) to 70% full signal (top gradient at right of panel: orange). Hierarchical clustering of antibodies by cross competition into 7 groups (plus a singleton labeled S2-3), enclosed in square boxes, with designations shown and in colors from dark blue (NTD-1) to dark red (S2-3). Green arrows on the left designate antibodies newly reported here. The lower parts of the panel show: competition of blocking antibody with soluble, human ACE2 (second gradient at right: dark red); log(IC_50_) in pseudovirus neutralization assay (third gradient at right: violet); area under the curve for ELISA binding (bottom gradient at right: brown); binding (ELISA) to recombinant domains and heterologous spike proteins. (**B**) Competition in cell-based assay for 36 antibodies with binding in ELISA format too weak for reliable blocking measurement (rows). Blocking antibodies (columns) selected from each of the 7 clusters in the ELISA assay (fig. S2). Strength of blocking shown as intensity of orange color, as in (A). (**C**) Distribution of antibodies from three individual subjects (expressed as percent of sequence pairs recovered from that subject) into the 7 principal clusters, plus a non-assigned (unknown) category (unk) and S2-3. Data are shown for only those subjects from whom we recovered at least 10 heavy-and light-chain sequence pairs. Heat map scale shown at right of panel. Top row shows total distribution, from panel (A) and (B).

Cell-based competition showed that mAbs with affinities too low to test by ELISA mapped primarily to S2 and to the NTD (Figs 2B and S3C). Results from including ELISA-mapped antibodies in the cell-based competition assay showed that the two assays were consistent (fig. S5A and B) and suggested that cell-surface binding simply extended the dynamic range of the ELISA competition assay to include less tightly binding antibodies, justifying use of the combined competition results in subsequent analyses. This combined approach showed that frequencies of cluster-targeting mAbs from the two individuals that contributed the most clones (C12 and G32) were largely similar to all others (Fig. 2C).

### Structural features of competition groups

We included in the competition assays, antibodies for which published structures show their interaction with S. We also determined by cryo-EM structures of Fab fragments of four mAbs from the RBD-1 and NTD-1 clusters bound with S ectodomain, to fill gaps in the representation of antibodies from those clusters in published work. Two of those structures are at relatively high resolution (those of Fab C12C9, in NTD-1, and Fab G32R7, in RBD-1), a third (C81C10, at the periphery of NTD-1) at intermediate resolution, and a fourth (12C11, in NTD-1) at much lower resolution.

*RBD-1.* The complex with Fab G32R7 (Fig. 3) has three RBDs in the “up” configuration, each bound with a Fab. The epitope is part of the RBD surface that faces outwards in the “down” configuration of the domain, but interference of the bound Fab with the NTD of the anticlockwise neighboring subunit (as viewed from “above” the spike in the orientation generally shown) would prevent binding to a down-oriented RBD. The connection of the RBD allows a range of orientations for the up configuration, and association with the G32R7 Fab does not fix the orientation of the RBD, blurring density in a 3-D reconstruction of the intact spike. Local refinement of an RBD-Fab subparticle then yielded a map that allowed us to build a good model of the interface (Fig. 3 and figs. S6, S7). RBD contacts are all with the heavy chain variable domain (V_H_), principally CDRH2, framework residues in the C”, D and E strands, and CDRH3. The unusually long CDRH3 (24 residues) also interacts with three glycans -- one on RBD Asn343 and the others on NTD Asn122 and Asn165. Although V_H_ approaches the NTD closely enough to interact with the glycans, we could identify just one likely additional contact with an NTD side-chain (Phe157). The one published RBD-1 antibody structure is that of S309, a neutralizing antibody isolated from a convalescent SARS-CoV donor that also neutralizes SARS-CoV-2 (*20*). Its contacts with the RBD do not overlap those of G32R7, but the light chain of the latter would collide with it.

**Fig. 3.**
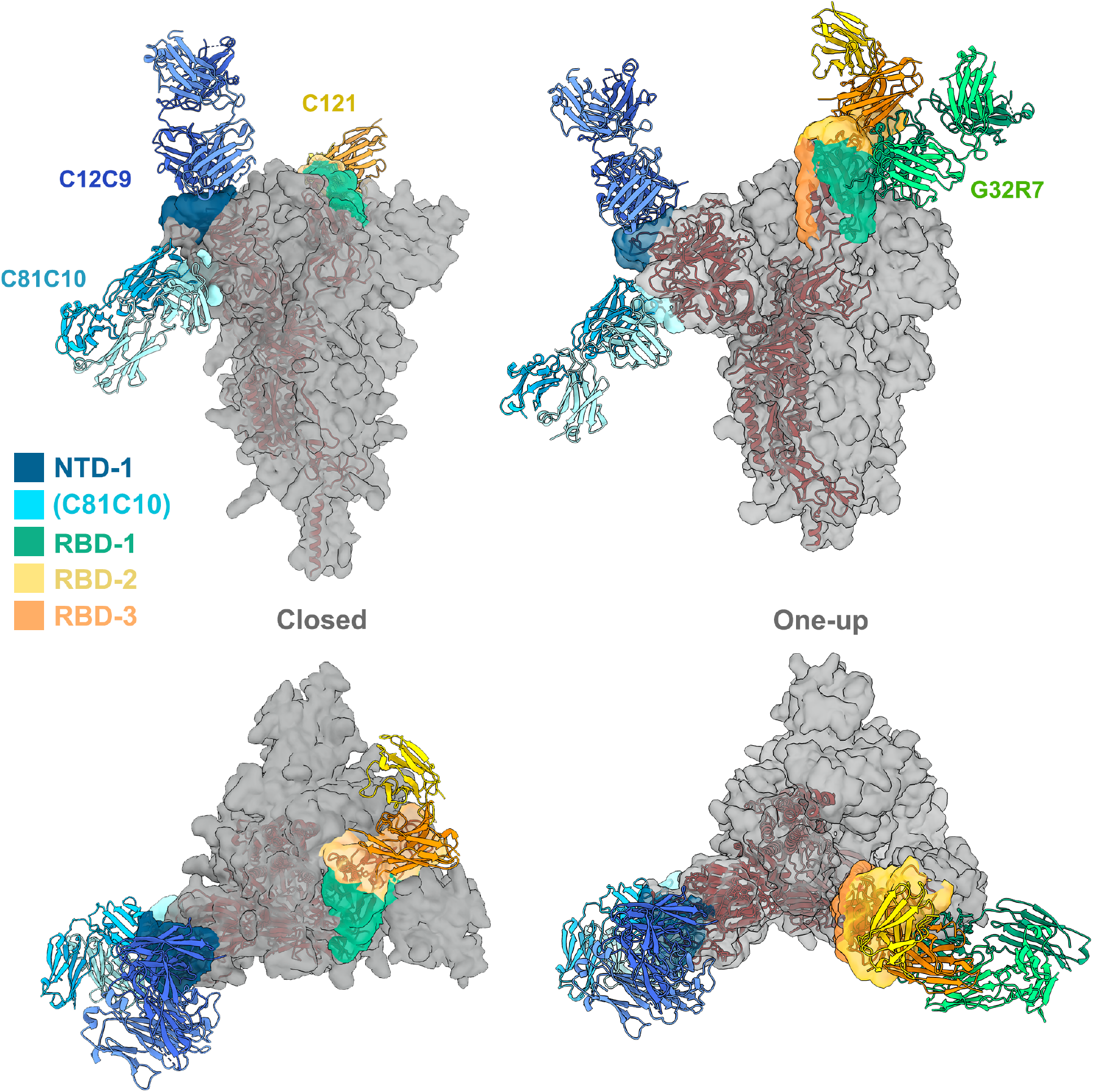
Ab contact regions. Surface regions of the SARS-CoV-2 spike protein trimer contacted by antibodies in four of the seven principal clusters, according to the color scheme shown (taken from the color scheme in Fig. 2), with a representative Fab for all except RBD-3. The C81C10 Fab defines an epitope just outside the margin of NTD-1, but it does not compete with any antibodies in RBD-2. The RBD-2 Fv shown is that of C121 (PDB ID: 7K8X: Barnes et al, 2020), which fits most closely, of the many published RBD-2 antibodies, into our low-resolution map for C12A2. Left: views normal to and along threefold axis of the closed, all-RBD-down conformation; right: similar views of the one-RBD-up conformation. C121 (RBD-2) can bind both RBD down and RBD up; G32R7 (RBD-1) binds only the “up” conformation of the RBD. The epitopes of the several published RBD-3 antibodies are partly occluded in both closed and open conformations of the RBD; none are shown here as cartoons. A cartoon of the polypeptide chain of a single subunit (dark red) is shown within the surface contour for a spike trimer (gray).

*RBD-2 and RBD-3.* Most potently neutralizing antibodies cluster in RBD-2; many published structures show modes of antibody binding within this group (*25, 26*). Their epitopes include various parts of the ACE2 binding site (i.e. the RBM) at one apex of the domain. For the antibodies characterized here, low resolution structure of Fab C12A2 showed that its epitope was essentially identical to that of published antibody 2-4 (*12*). The same IGVH encodes the heavy chains of both antibodies, and the light-chain genes are closely related (overall amino-acid sequence identity). They contact the slight concavity in the center of the RBM, the site for most of the neutralizing antibodies represented by structures in the PDB. The probably immunosubdominant RBD3 class includes several antibodies for which published structures are available; we included CR3022, an antibody originally isolated from a

SARS-CoV convalescent subject that cross-reacts with SARS-CoV-2 (*27*). Its epitope on the RBD is nearly opposite that of G32R7 (Fig. 3), in an epitopic region partly occluded in the down configuration of the RBD previously referred to as a “cryptic supersite” (*26*).

*NTD-1.* NTD-1-cluster antibodies vary in neutralizing strength from strong (e.g., C12C9) to weak (C12C11). The latter, judging from the low-resolution map (fig. S8) appears to have a footprint that coincides with that of the published 4A8 antibody (PDB ID: 7C2L; (*22*)). Like the G32R7 complex, the C12C9 complex also required local subparticle refinement to yield a map interpretable at the level of side-chain contacts at the Fab-NTD interface. Its footprint overlaps that of 4A8, but it is displaced slightly toward the threefold axis of the S trimer. Both antibodies have principal contacts in two NTD surface loops, residues 140-160 and 245-260. The C81C10 mAb, which we have grouped in NTD-1 but which competes with only two of the most weakly binding members of that cluster, appears to represent a distinct and potentially subdominant subset. An 8 Å resolution structure (fig. S8) bound with spike trimer shows that its epitope is at the “bottom’ of the NTD, well displaced from the epitopes of C12C9, C12C11 and 4A8.

*NTD-2.* We have so far no structures of NTD-2 antibodies bound with spike, but from noncompetition with NTD-1, insensitivity to NTD loop deletions (see below), and exposure of NTD surfaces on the trimer, we suggest that the NTD-2 epitopes may be on one of the lateral faces of the NTD.

### Representation of neutralizing antibodies in RBD and NTD clusters

Using two different pseudovirus assays, we determined neutralization by mAbs from each of the 7 clusters and found neutralizing antibodies in 5 of the 7 clusters (RBD-1, -2, -3, NTD-1, and NTD-2) (fig. S9). The most potent were in RBD-2, as expected from their co-clustering with known strong neutralizers such as REGN10987 and REGN10933, which are used as a mAb drug cocktail (*28*) and from many previous reports (*12, 13, 29*). Among Abs from that cluster, 52-58% neutralized with IC_50_<0.1 μg/ml and 28.5-35% with IC_80_<0.1 μg/ml (fig. S9D). The strongest of the RBD-1 antibodies had IC_50_ in the range of 1 μg/ml; those in RBD-3 were in general much weaker. NTD-1 antibodies appeared more sensitive to the neutralization assay used, but a few, such as C12C9, approached or exceeded the strongest in RBD-1 (fig. S9A). NTD-2 antibodies were in general less potent than NTD-1 antibodies, and none of the S2 antibodies neutralized infection, with the possible exception of very weak neutralization by G32C5 (IC_50_ of 22 ug/ml) (fig. S9C).

### Molecular features of mAb recognition groups

Variable region exons of IgH and IgL genes are each assembled by V(D)J recombination from a diversity of gene segments. Preference of VH gene segment usage frequencies differed among the 7 mAb clusters (Fig. 4A). Enrichment for VH3-53, previously reported to be associated with SARS-CoV-2 S (*21*), was exclusively within the RBD-2 group. VH3 family antibodies are particularly abundant in all the clusters. VH3-30 and VH3-30-3, which have average frequency in the general human repertoire of 5.4% and 1.3% respectively, account between them for over 30% of the antibodies in RBD-1, and for 16 of the 19 antibodies in S2-2. The VH1 and VH4 families are co-dominant with VH3 in NTD-1 and NTD-2, respectively (Fig. 4A). VH1-69 encoded antibodies are enriched in S2-1, which contains most of the cross-reactive antibodies to other coronaviruses. VH1-69 encoded antibodies are frequently observed in antiviral responses to influenza virus, HCV, and HIV-1 (*30*), and previous work reported that SARS-CoV-2 spike-specific mAbs isolated from SARS-CoV infected patients also showed an enriched VH1-69 gene segment usage (*31*). VH1-69, which is well represented in heavy chains of “natural antibodies”, also associates strongly with polyreactivity. VH and VL somatic mutation levels were generally, but not significantly, greater in S2-1 (fig. S10A)

**Fig. 4.**
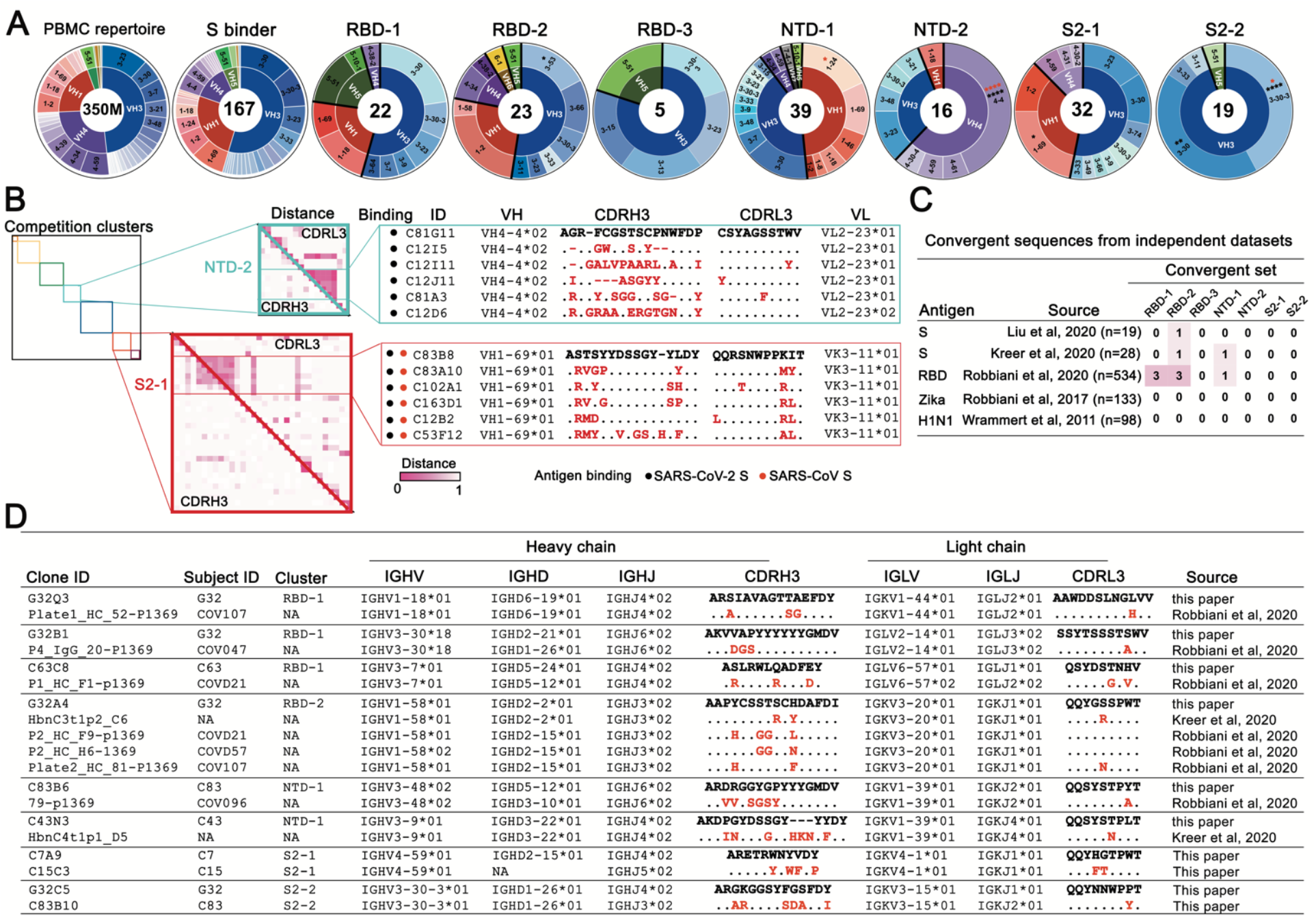
Antibody sequence analyses. **(A)** Heavy-chain variable-domain genes of the 167 mAbs characterized by binding SARS-CoV-2 spike in either ELISA or cell-surface expression format. The inner ring of each pie chart shows the VH family and the outer ring, the gene. PBMC repertoire is from 350 million reads of deep sequencing (*37*). S binders include 167 clones in Table S2. *\*P* < 0.05, *\*\*P* < 0.01, \*\*\**P* < 0.001, \*\*\*\**P* < 0.0001; Bonferroni correction. Red asterisks: comparing to S binders; black asterisks: comparing to a non-selected B cell repertoire from PBMCs. **(B)** Maps of pairwise distances of CDRH3 (lower left triangle) and CDRL3 (upper right triangle) for the NTD-2 and S2-1 cluster antibodies from (A). Antibodies in both clusters arranged by VH usage. Clones converging on identical VH/VL alleles and closest distance of CDRL3 from the same cluster are shown. Pairwise distances analyzed by Mega X. Intensity of color shows the distance, from 0 (identical) to 1 (no identity). Sequence alignment for the antibodies from the indicated clusters with identical VH and VJ and similar CDR3s. Differences in CDR3s from the reference sequence (bold) are in red; dashes indicate missing amino acids; dots represent identical amino acids. **(C)** Summary of convergent sequences of anti-SARS-CoV-2 S and RBD antibodies from independent datasets. Ig sequences derived from binding to DIII of Zika virus E protein, and HA of influenza virus H1N1 were used as control datasets. Convergent sequences had identical VH and VL and >50% identity in CDRH3 and CDRL3. **(D)** Representative convergent clones from different individuals and independent datasets from Fig. 4C.

IgH and IgL variable regions harbor three complementary determining regions (CDRs), which serve as principal contact sites for antigen. CDRs 1 and 2 for H and L chain are encoded within the VH and VL gene segments, respectively. Highly diverse, non-templated sequences produced by VDJ_H_ junctions encode CDRH3 regions, which have dominant roles in most Ab-antigen interactions. CDRL3, which can contribute antigen contact surfaces, is also diverse due to VJ_L_ junctional heterogeneity, but has less non-templated sequence additions. Intracluster mAb CDR3 sequence comparisons showed little sequence similarity (fig. S10B). NTD-2 contained a subcluster of identical CDRL3 sequences that were associated with the same VH and VL segments from two different individuals (C81, C12) (Fig. 4B). S2-1 had a small subcluster of CDRH3 and CDRL3 sequence similarities from 5 different study participants (C83, C102, C163, C12, C53) (Fig. 4B). These data indicate substantial intracluster CDR3 diversity with rare instances of CDR3 sequence similarity between different individuals.

We also asked whether we could find sequences very similar (i.e. convergent) to any in our dataset, from other COVID-19 data sets for which paired IgH and IgL sequence data are available. Based on prior convergent sequence analysis (*32*), our criteria for convergence were (i) same VH and VL, and (ii) no less than 50% CDRH3 and CDRL3 identity, These criteria identified rare sequences very similar to representatives from RBD-1, RBD-2, and NTD-1 in independent datasets for SARS-CoV-2 (*1, 12, 33*), but not for antibodies against Zika (*34*) or influenza (*35*) viruses (Fig. 4C, D). We also found convergent pairs within our own dataset representing both S2-1 and S2-2 (Fig. 4D).

Finally, we note that antibody C93D9 represents a striking example of structural convergence. All but two of the 20 antibodies from the literature shown in fig. S11C have the same VH and a non-random selection of VL but divergent CDRH3 sequences and lengths. Nonetheless, all 20, as well as C93D9, bind the RBM in almost identical poses—consistent with germline encoded CDRs as the principal binding contacts (*21*).

### Reactivity of the anti-S memory B cell repertoire for emerging SARS-CoV-2 variants

Emergence of SARS CoV-2 variants that enhance transmissibility, such as the variant B.1.1.7 (i.e. the UK variant) (*36*), and in some cases reduce the neutralization titers of convalescent sera, such as the variant B.1.351 (i.e. South Africa (SA) variant) (*37*), indicates more rapid evolution of the virus than expected from the error-correcting properties of coronavirus RNA-dependent RNA polymerases. In the case of the SA variant in particular, the clusters of three substitutions and one deletion in the NTD and three substitutions in the RBD concentrate at contacts of the most potent of the many well-characterized neutralizing antibodies. Moreover, recurrent deletions in loops of the NTD appear to accelerate SARS CoV-2 antigenic evolution (*38*).

We examined the effects of naturally occurring mutant spike protein on binding of mAbs in each competition group. The UK variant had lower affinity for various mAbs in the RBD-1 and NTD-1 cluster. None of the RBD-1 mAbs lost binding completely, and testing a variant with just the deletion at position 144 in the NTD showed that this single mutation caused loss of binding by nearly two-thirds of the mAbs in the NTD-1 cluster. Mutations in the variant B.1.351 had more pronounced effects, particularly on mAbs in the RBD-2 and NTD-1 clusters, as expected from the positions of the sequence changes. In addition to the N501Y substitution also present in the variant B.1.1.7, an E484K mutation lies at the center of the epitope for many of the most potent RBD-2 neutralizing antibodies. About one-third of the RBD-2 mAbs retained modest to high affinity, but the variant spike failed to bind any of the NTD-1 cluster, with the marginal exception of 4A8.

Differential effects on antibodies with overlapping but still distinct epitopes illustrate the potential importance of a redundant, polyclonal response. Although C12C9, C12C11, and 4A8 all contact the 140-160 loop (Fig. 3 and S7) and all are sensitive to the multi-position, Δ141-144 and Δ243-244, recurrent deletions, only the latter two are sensitive to the recurrent, single-position deletions at 144 or 146 (Fig. 5 and fig. S12). Moreover, although they are in the same convergent structural class whose members bind the RBM in nearly identical poses (fig. S11), CC12.1 fails to recognize B.1.351 (~0%) while C93D9 retains some marginal affinity (~27%) (Fig. 5). Thus, apparently redundant memory B cell clones can have non-redundant functional roles.

**Fig. 5.**
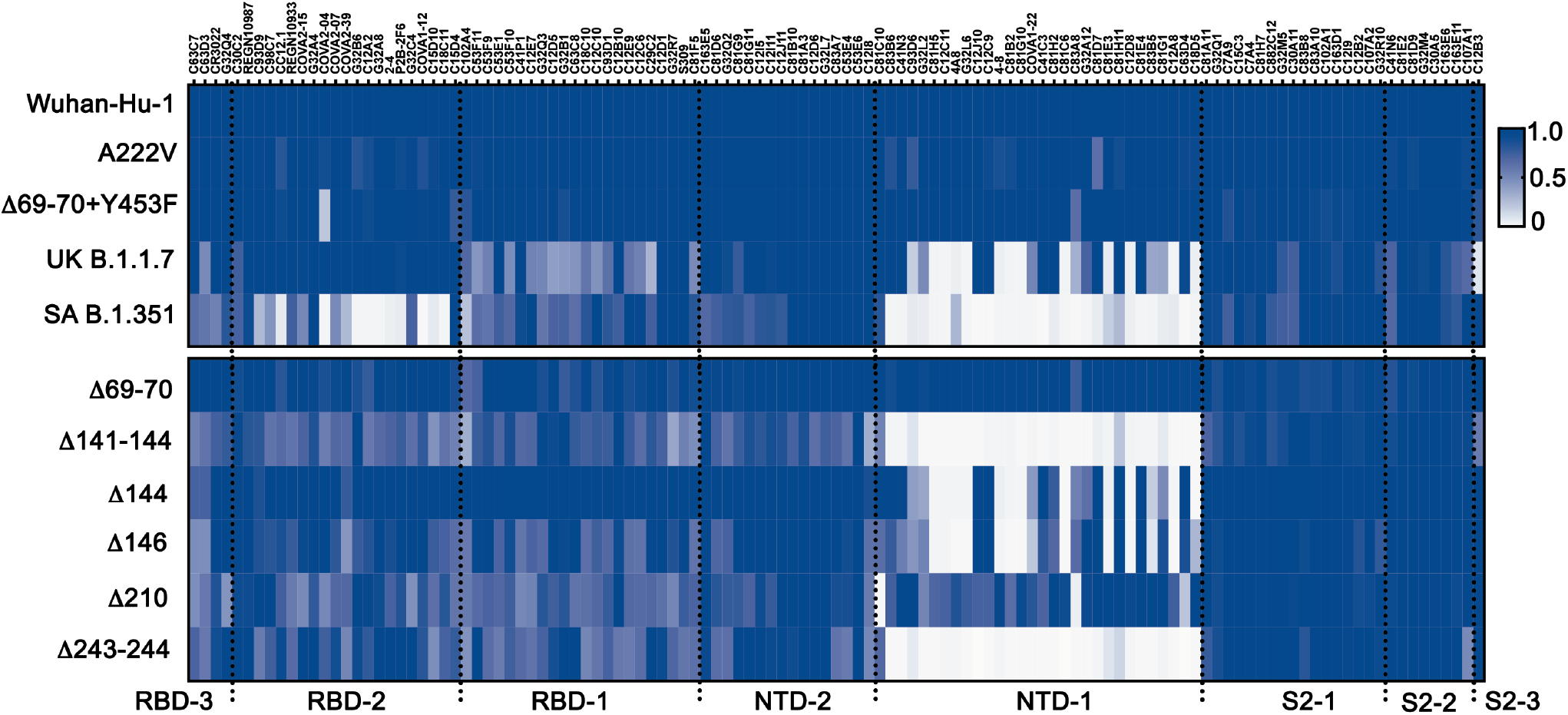
Recognition of naturally occurring deletions and mutations in the spike. Heat map showing binding of 119 mAbs to Nextstrain cluster 20A.EU1 (A222V), Danish mink variant (Δ69-70 and Y453F), UK B.1.1.7 (Δ69-70, Δ144, N501Y, A570D, P681H, T716I, S982A, D1118H) and SA B.1.351 (L18F, D80A, D215G, Δ242-244, K417N, E484K, N501Y, A701V) (top) and NTD deletion variants (bottom). The Wuhan-Hu-1 S sequence and all variants include the D614G mutation. Binding for each mAb was first normalized (“normalized IgG MFI”) by dividing the MFI for that mAb by the MFI for C81E2 (S2-2 cluster). The normalized MFI of for binding the Wuhan-Hu-1 spike was used as a reference (normalized Wuhan IgG MFI). The relative binding intensities of the tested mAbs for each variant, calculated as the ratio of the normalized variant IgG MFI and the normalized Wuhan IgG MFI, are shown in shades of blue.

## DISCUSSION

Our results illustrate the landscape of memory B cell coverage of the SARS-CoV-2 S glycoprotein in convalescent donors. Unlike the terminally differentiated plasma cells that determine the profile of serum antibodies, memory B cells will clonally expand upon re-exposure to antigen, some differentiating into fresh antibody secreting cells and others re-entering germinal centers and undergoing further SHM-mediated diversification and affinity maturation. These outcomes offer a layer of flexibility for adaptation to drifted or related viral strains, if available secreted antibodies fail to prevent initial infection. Loss of protection against overt or severe disease is not an inevitable consequence of a waning serum antibody titer. This atlas of B cell memory therefore maps systematically a crucial component of the long-term immune response to SARS-CoV-2 infection.

The donors for this study experienced COVID-19 symptom onset between March 3 and April 1, 2020, and blood draws analyzed here were between April 2 and May 13, 2020, early in the pandemic. Immune responses in these SARS-CoV-2 naive donors were to early and relatively homogeneous variants circulating well before emergence of the UK and SA strains first reported in Dec. 2020 and probably before the spread, in New England, of the D614G variant that in any case did not substantially alter antigenicity (*39, 40*). This set of BCR sequences and corresponding mAbs thus represents responses to a relatively homogeneous infectious virus and provides a valuable tool for examining the degree to which these antibodies retain recognition of emerging variants and for studying the extent to which loss of neutralizing titer correlates with loss of longer-term protection.

The competition clusters we have identified are roughly analogous to genetic complementation groups. Competition can result from overlapping binding footprints or non-overlapping but neighboring footprints that lead to mutual exclusion of IgGs bound at the two adjacent epitopes. Competition can also result from stabilization by one antibody of a conformation (e.g., the up-down conformational isomerism of the RBD) that excludes or lowers affinity of the another. Any of these mechanisms may contribute to the clusters we have mapped, but the outcome in all cases is an apparent redundancy of binding capacity in a broadly polyclonal response that may nonetheless impart recognition breadth toward an evolving pathogen within a single individual.

Complementary recognition of non-overlapping viral targets by non-competing antibodies in the repertoire can reduce the likelihood of viral escape (*41*). Our data suggest an additional mechanism for preventing viral escape: competing antibodies may help retain recognition of a rapidly evolving antigen by their differential sensitivity to specific mutations. The potential dynamic reach of otherwise redundant mAb recognition, illustrated by selective retention of affinity for the UK variant by some antibodies within a cluster but not by others, may give selective advantage to immune mechanisms that yield multiple competing antibodies to critical epitopes, as those that retain adequate affinity can then re-activate, expand, and potentially undergo further affinity maturation. The emergence of strains that may have gained selective advantage by escape from neutralization emphasizes the importance of determining whether the level of retained affinity for the S protein by some antibodies in the immunodominant clusters influences protection from clinical disease.

## Acknowledgments

We thank the study volunteers, Sudeshna Fisch and Reem Abbaker for support in patient recruitment and sample collection, and Losyev Grigoriy for flow cytometry cell sorting. We thank Tianshu Xiao for providing human ACE2 protein and Ning (Alexa) Guan for manuscript review. This study was supported by National Institutes of Health grants T32 AI007245 (to J.F.), T32 GM007753 (to B.M.H.), AI146779 (to A.G.S.), AI007512 (to A.Z.), T32 AI007306 (to Y. Chen) and AI121394, AI139538, and AI137940 (to D.R.W.). W.P.D and K.R.M were supported by the University of Pittsburgh, the Center for Vaccine Research, and W.P.D was supported by The Richard King Mellon Foundation, the Henry L. Hillman Foundation and the Commonwealth of Pennsylvania, Department of Community and Economic Development. Work in the laboratories of A.G.S., B.C., S.C.H. and D.R.W were funded by the Massachusetts Consortium on Pathogenesis Readiness. D.R.W. acknowledges support from the Food Allergy Science Initiative, the Massachusetts Institute of Technology Center for Microbiome Informatics and Therapeutics, and Fast Grant funding for COVID-19 science. D.R.W also acknowledges support from the Ragon Institute, and the Mark and Lisa Schwartz and the Schwartz Family Foundation and acknowledges the interest of Enid Schwartz. Support for the Harvard Cryo-EM Center for Structural Biology came from the Nancy Lurie Marks Family Foundation.

## Author contributions

D.R.W. designed the study. P.T. and A.G. conducted ELISA and FACS experiments. I.W.W. and G.B. conducted cryo-EM experiments. P.T., M.T., A.Gautam., Y. Chen, A.Z., T.Z., and J.L. performed single cell sorting and antibody cloning. P.T. and J.F. prepared yeast expression constructs. P.T., K.M., W.P.D., L.J.R., K.R.M., and Y.Cai prepared spike variant expression constructs. P.T., A.Gautam., I.W.W., G.B., N.G., N.B.W., S.C.H. and D.R.W analyzed data. M.T., F.J.N.L., and A.Gautam. recruited patients and processed samples. P.T., C.L.L., M.T. and M.S.S. performed pseudovirus neutralization assays and analysis. L.G.A.M. and A.Griffiths. performed authentic virus neutralization assays and analysis. M.T., I.W., E.A.M., G.B., J.F., B.M.H., A.G.S., Y.Cai, and B.C. contributed recombinant protein. N.G., N.B.W. P.T. prepared the first draft. S.C.H. and D.R.W. supervised research and wrote the paper.

## Data and materials availability

The cryo-EM maps and atomic models will be deposited at the EMDB and PDB. Sequences of the monoclonal antibodies characterized here will be deposited at the GenBank. Reagents and materials presented in this study are available upon request, in some cases after completion of a materials transfer agreement.

## Supplementary Information

Materials and Methods

Figs. S1 to S12

Table S1

References (*42–52*)

Data S1 to S2

## SUPPLEMENTARY INFORMATION

### Materials and Methods

#### Human samples

The study and protocol were approved by Partners Institutional Review Board. Volunteers aged 18 and older with a history of COVID-19 were enrolled between March and May 2020. COVID-19 was diagnosed by a healthcare professional based on symptoms and a positive nasopharyngeal swab RT-PCR test except for G32, who was diagnosed by an antibody test. Participants self-reported data for body-mass index (BMI), symptom onset and recovery dates and self-rated the severity of their COVID-19 symptoms on a 1-10 scale, with 1 describing very mild symptoms and 10 describing very severe symptoms. Blood samples were collected at least 2 weeks after symptom resolution. Symptom duration is the time between symptom onset and recovery dates. Detailed information about the cohort is in Data S1. BMI was calculated as participant’s weight in kilograms divided by the square of their height in meters. Blood samples were processed within 4 hours of sample collection. PBMCs and plasma samples were isolated by density gradient centrifugation with Ficoll-Paque PLUS (GE Healthcare) and stored at −80°C until use.

#### SARS-CoV-2 spike-specific single B cell sorting

B cells, enriched from PBMCs with human CD19 MicroBeads (Miltenyi), were incubated with 2 ug/ ml flag-tagged S protein or mixture of flag-tagged S protein (Genscript, Cat. Z03481) and His-tagged RBD (*10*) on ice for 30 minutes. Cells were then washed with 2% FBS (Hyclone) in PBS and stained with mixture of anti-human IgG (Percpcy5.5; Biolegend Cat. 410710), anti-human IgD (FITC; Biolegend Cat. 348205), anti-human IgM (Bv605; Biolegend Cat. 314524), anti-CD27 (APCcy7; Biolegend Cat. 356404). A mixture of PE- and APC-conjugated anti-flag antibodies (Biolegend Cat. 637309 and 637307) was also added for gating S-specific double positive cells, or a mixture of PE-conjugated anti-His (Biolegend Cat. 362603) and APC-conjugated anti-flag, for S positive and RBD negative cells. Memory B-cells were gated on DAPI’CD19^+^IgM’IgD’IgG^+^CD27^+^. Individual S double positive or S+RBD-cells were sorted with a FACSAria Fusion (BD Biosciences) into each well of 96-well microplates containing 4 μl/well of lysis buffer (0.5X PBS, 10 mM dithiothreitol, and 4U RNaseOUT). Lysed cells were immediately frozen and stored at −80 °C until use.

#### Antibody cloning and production

Cloning and expression of mAbs from single, SARS-CoV-2 S-specific B-cells were performed as described previously (*10*). In brief, mRNA from lysed B cells was reverse transcribed with SuperScript III (ThermoFisher) and random hexamers. Two rounds of PCR were performed to amplify heavy and light chain transcripts. Amplified products from the second round PCR were detected by agarose gel and further verified by Sanger sequencing. Sequences were analyzed with IgBlast (https://www.ncbi.nlm.nih.gov/igblast/), and sequence confirmed PCR products were then amplified with gene specific primers containing restriction enzyme sites for cloning into human IgG1, kappa and lambda expression vectors (gifts from Michel C. Nussenzweig, Rockefeller University). For small scale antibody production, paired heavy and light chains were co-transfected into HEK 293T cells (ATCC, Cat. CRL-3216) in 6-well plates with Lipofectamine 3000 (ThermoFisher, Cat. L3000015) following manufacturer’s instructions. Cells were cultured in DMEM supplemented with 10% fetal bovine serum (FBS) and incubated at 37°C with 5% CO_2_. The medium was replaced with fresh medium at 12 hours post-transfection and the supernatant harvested after 48 hours. Cell debris were removed by centrifugation at 2000 g for 10 min and the cleared supernatant stored at 4°C for further use.

For large scale antibody production, paired heavy chain and light chain were co-transfected in Expi293F cells (ThermoFisher, Cat. A14527) with ExpiFectamine (ThermoFisher, Cat. A14525) in 250 ml Erlenmeyer flasks following manufacturer’s instructions. Cells were cultured in Expi293 expression medium at 37°C and 8% CO_2_ with shaking at 125 RPM. On day 7, cells were removed by centrifuging at 2000 RPM for 10 min. Clear supernatants were incubated overnight at 4°C with protein A agarose beads (ThermoFisher, Cat. 20334), followed by washing with PBS, elution with 0.1 M Glycine (pH 2.7) and neutralization with 1 M Tris-HCl (pH 8.0). Purified antibodies were dialyzed against PBS for further use.

#### Monoclonal antibody screening with ELISA

SARS-CoV-2 S protein and the RBD proteins of other coronaviruses were prepared as described (*10*). SARS-CoV-2 S2 and NTD proteins were purchased from Sino biological (PA, USA). ELISA was carried out as described (*10*). Briefly, MaxiSorp 96-well ELISA plates (ThermoFisher) were coated with 50 ng/well of the antigen in PBS at 4°C overnight. Plates were blocked with 150 μl of 4% BSA in PBS for at least 2 hours. Supernatant of antibody-expressing cells was diluted 10-fold for the first well and 4-fold serial diluted for subsequent wells, applied to the plates, and incubated at 4°C overnight. Plates were washed 4 times with PBS supplemented with 0.05% Tween-20 (PBST). Anti-human IgG-alkaline phosphatase (Southern Biotech, Birmingham, AL) at a final concentration of 1μg/ml in 1% BSA and 0.05% Tween-20 was added at 50 μl/well and incubated for 1 hour at room temperature. Plates were washed three times with PBST. Developing solution (0.1 M Glycine, pH 10.4, with 1 mM MgCl_2_ and 1 mM ZnCl_2_, containing alkaline phosphatase substrate p-nitrophenyl phosphate (Sigma-Aldrich) at final concentration of 1.6 mg/mL) was then added to the plates at 100 μl/well, and incubated for 2 hours at room temperature. Absorbance was measured at 405 nm by microplate reader (Biotek Synergy H1).

For half maximal effective concentration (EC_50_) and area under curve (AUC) analysis of antibody binding by ELISA, 5-fold serial dilutions were produced from an initial concentration of 10 μg/ml of purified antibody in the ELISA procedure described above. EC_50_ and AUC were calculated with GraphPad Prism 9.

#### Cell surface binding assays

Antibodies were tested for binding to surface-expressed SARS-CoV-2 spike, RBD, and NTD on HEK 293 T cells (spike) and on yeast (RBD, NTD). For yeast expressing, RBD and, RBD (aa 319-529) and NTD (aa 17-286) were cloned into a pCHA vector (gift of K. Dane Wittrup, Koch Institute for Integrative Cancer Research, MIT, Cambridge, MA). Plasmids were chemically transformed into yeast cell line EBY100 to display RBD or NTD as previously described (*10*). Briefly, single clones were cultured in SDCAA selection medium for 48 hours at 30 °C and 250 RPM. Cells were pelleted and resuspended in SGCAA medium to an absorbance of 0.5-1 at 600 nm, cultured at 20°C with shaking at 250 RPM for another 48 hours to induce expression. RBD and NTD were detected by anti-c-Myc IgY antibody (ThermoFisher, Cat. A21281). Yeast expressing RBD or NTD were incubated with antibody supernatant and anti-c-Myc IgY on ice for 30 minutes and then washed with PBS with 2% FBS twice. Cells were then stained with goat anti-chicken IgG (Alexa Fluor 488, ThermoFisher, A11039) and goat anti-human IgG (Alexa Fluro 647, ThermoFisher, A21445) on ice for 15 min, followed by washing twice with PBS supplemented with 2% FBS (FACS buffer). Cells were resuspended in FACS buffer and detected by flow cytometry (BD Canto II). Data were analyzed by FlowJo 10.7.1. For analysis of antibody binding to SARS-CoV-2 S on HEK 293 T cells, a plasmid containing Wuhan-Hu-1 S (HDM-SARS2-spike-delta21, Addgene, Cat. 155130) was co-transfected with pmaxGFP (Lonza) in HEK 293T cells using Lipofectamine 3000. Fresh medium was added at 24 hours, and cells were harvested at 48 hours post-transfection in PBS with 2 mM EDTA. Cells were stained with antibody supernatant on ice for 1 hour, washed twice with FACS buffer, and stained with goat anti-human IgG (Alexa Fluro 647 ThermoFisher, A21445) and DAPi (to distinguish dead and live cells). After washing twice with FACS buffer, cells were resuspended in FACS buffer and detected by flow cytometry (BD Canto II). S^+^ cells were identified by gating on DAPi^-^GFP^+^. Data were analyzed in FlowJo 10.7.1. For half maximal effective concentration (EC_50_) of antibody binding to SARS-CoV-2 S in the cell-based assay, eight three-fold serial dilutions of purified antibody were produced starting from 10 μg/ml, followed by flow cytometric binding analysis as above. EC_50_ was calculated with GraphPad Prism 9.

#### ELISA-based antibody competition

The competition assay was performed as described (*12*). Briefly, detection antibodies were biotinylated with EZ-Link™ Sulfo-NHS-LC-Biotin (ThermoFisher) according to manufacturer’s protocol. 50 ng/well of SARS-CoV-2 S protein were coated on ELISA plates at 4 °C overnight. Plates were blocked with 150 ul of 4% BSA in PBS for 2 hours. 30 μl of 2 μg/ml biotinylated antibody were mixed with 30 μl of 200 μg/ml blocking antibody and added to ELISA plates. For antibody competition with hACE2 (aa18-615), 100 ng/well hACE2 were coated on ELISA plates at 4 °C overnight. Plates were blocked with 4% BSA, and a mixture of 30 μl of 2 μg/ml Twin-Strep-tag HexaPro S (*42*) and 30 μl of 200 μg/ml blocking antibody was then added. Plates were incubated for 2 hrs at 37°C and washed 4 times with PBST. 50 μl/well of streptavidin-alkaline phosphatase (BD Biosciences, Cat. 554065) was added to the wells using a dilution of 1:1000 dilution of the stock solution according to the manufacturer’s instructions, and incubated for 1 hour at room temperature. Plates were washed 5 times with PBST and developed at room temperature for 2 hours. Absorbance was measured at 405 nm by microplate reader (Biotek Synergy H1). The detection signal was calculated by (OD value of mixture antibodies-OD value of PBS)/ (OD value of biotinylated antibody alone-OD value of PBS) x100%. Negative values were treated as 100% competition.

#### Cell-based antibody competition

S (HDM-SARS2-spike-delta21, Addgene, Cat. 155130) and GFP (pmaxGFP) was co-expressed in HEK 293T cells. 30 μl of 2 μg/ml biotinylated antibody were mixed with 30 μl of 200 μg/ml blocking antibody and added to cells. After 1-hour incubation on ice, followed by washing twice with FACS buffer, cells were stained with 50 μl of 1:1000 diluted DyLight 649 Streptavidin (BioLegend, Cat. 405224) and DAPi. After washing twice with FACS buffer, cells were resuspended in FACS buffer and detected by flow cytometry (BD Canto II). S+ cells were gated on DAPi-GFP+. Data were analyzed by FlowJo 10.7.1. The detection signal was calculated by (MFI of mixture antibodies-MFI of PBS)/ (MFI of biotinylated antibody alone-MFI of PBS) x100%. Negative values were treated as 100% competition.

Antibody binding to S variants Variants included: Wuhan-Hu-1 S (HDM-SARS2-spike-delta21-D614G, Addgene, Cat. 158762), recurring NTD deletions as described (*38*) (Δ69-70, Δ141-144, Δ144, Δ146, Δ210, Δ243-24), Nextstrain cluster 20A.EU1 (A222V), Danish mink variant (Δ 69-70 and Y453F), UK B.1.1.7 (Δ69-70, Δ144, N501Y, A570D, P681H, T716I, S982A, D1118H) and SA B.1.351 (L18F, D80A, D215G, Δ242-244, K417N, E484K, N501Y, A701V). We note that plasmids with Δ144 and Δ145 have the same coding sequence due to the presence of tyrosine at both sites. All variants contain D614G. For binding, 10 μg/ml of antibody were incubated with cells, with goat anti-human IgG as secondary antibody for detection by flow cytometry (BD Canto II). S+ cells were gated on DAPi-GFP+. Data were analyzed by FlowJo 10.7.1. Antibodies were first normalized for each spike variant (normalized IgG MFI), by dividing MFI of tested mAb by MFI of C81E2 reactive to the S2 region. Normalized MFI of Wuhan-Hu-1 S was used as a reference (normalized Wuhan IgG MFI). Relative binding intensity of the tested mAb for different variants was calculated as the ration of normalized variant IgG MFI to normalized Wuhan IgG MFI. Relative binding signal > 1 was treated as no loss of binding and set to 1.

#### Pseudovirus production and neutralization assay

Pseudovirus particles were produced as described (*43*). HEK 293T cells were co-transfected with spike envelope plasmid (HDM-SARS2-spike-Δ21, Addgene, Cat. 155130), package plasmid (psPAX2, Addgene, Cat. 12260) and backbone plasmid (pLenti CMV Puro LUC, Addgene, Cat. 17477) with Lipofectamine 3000. Medium was replaced with fresh medium at 24 hours, and supernatants were harvested at 48 hours post-transfection and clarified by centrifugation at 300g for 10 min before aliquoting and storing at −80°C. SARS-Cov-2 pseudovirus neutralization assay was performed as described (*44*), with target cell line 293FT expressing human ACE2 and serine protease TMPRSS2 (provided by Marc C. Johnson, University of Missouri) or TZM.bl expressing human ACE2. Cells at 1.8 x 10^4^ cell/well were seeded in 96-well plates 16 hours in advance. Serial diluted mAb was mixed with pseudovirus and incubated for 1 hour at 37°C before adding to cells. Cells infected without mAb were scored as 100% infection; cells cultured without pseudovirus or mAb as blank controls. After 48 hours incubation at 37°C with 5% CO_2_, cells were processed with luminescent regent (ONE-Glo^TM^, Promega) according to manufacturer’s instructions, and luminescence (RLU) was measured with a microplate reader (Biotek Synergy H1). Inhibition was calculated by 100-(RLU of mAb-RLU of blank)/ (RLU of pseudovirus-RLU of blank) x100%. Values for half inhibition (IC_50_) and 80% inhibition (IC_80_) were calculated with GraphPad Prism 9.

#### Authentic virus neutralization assay

Cell Culture: NR-596 VeroE6 cells (BEI Resources) were maintained in Dulbecco’s modified Eagle medium (DMEM) (Gibco™) with 10% heat inactivated fetal bovine serum (Gibco™), GlutaMAX (Gibco™), non-essential amino acids (Gibco™) and sodium pyruvate (Gibco™). One day prior to the assay, VeroE6 cells were seeded at a density of 8.0 x 10^5^ cells per well of a 6-well plate (Falcon™ Polystyrene Microplates, Cat. 353934) in 2 mL media.

Virus propagation: Passage 4 SARS-CoV-2 USA-WA1/2020 was received from the University of Texas Medical Branch. A T225 flask of VeroE6 cells was inoculated with 90μL starting material in 15mL DMEM/2 % HI-FBS (Gibco™). The inoculated flasks were incubated in a humidified incubator at 37°C/5% CO_2_ with periodic rocking for 1 hour. After 1 hour, 60mL of DMEM/2% HI-FBS (Gibco™) was added without removing the inoculum and flasks were then incubated again at 37°C/5% CO_2_. Flasks were observed daily for progression of CPE and stock was harvested at 66 hours postinoculation. Stock supernatant was harvested and clarified by centrifugation at 5,250 RCF at 4°C for 10 minutes and heat inactivated fetal bovine serum concentration (Gibco™) was increased to a final concentration of 10%.

Viral neutralization reduction assays were performed at biosafety level 4 (BSL-4) at the National Emerging Infectious Disease Laboratories (NEIDL). An Avicel plaque reduction assay was used to quantify plaques. Antibody samples were serially diluted in Dulbecco’s Phosphate Buffered Saline (DPBS)(Gibco™) using two-fold dilutions. Dilutions were prepared in triplicate per antibody and plated in triplicate. Each dilution was incubated at 37 °C and 5% CO_2_ for 1 hour with 1000 plaque forming units/ml (PFU/ml) of SARS-CoV-2 (isolate USA-WA1/2020). Controls included DPBS as a negative control and 1000 PFU/ml SARS-CoV-2 incubated with DPBS. The maintenance medium was removed from each plate and 200 μL of each inoculum dilution was added to confluent monolayers of NR-596 Vero E6 cells (including a positive and mock negative control) in triplicate and incubated for 1 hour at 37°C/5% CO_2_ with gentle rocking every 10-15 minutes to prevent monolayer drying. The overlay was prepared by mixing by inversion Avicel 591 overlay (DuPont Nutrition & Biosciences, Wilmington, DE) and 2X Modified Eagle Medium (Temin’s modification, Gibco™) supplemented with 2X antibiotic-antimycotic (Gibco™), 2X GlutaMAX (Gibco™) and 10% fetal bovine serum (Gibco™) in a 1:1 ratio. After 1 hour, 2 mL of overlay was added to each well and the plates was incubated for 48 hours at 37°C/5% CO_2_. 6-well plates were then fixed using 10% neutral buffered formalin prior to removal from BSL-4 space. The fixed plates were then stained with 0.2% aqueous Gentian Violet (RICCA Chemicals, Arlington, TX) in 10% neutral buffered formalin for 30 min, followed by rinsing and plaque counting. The half maximal inhibitory concentrations (IC_50_) were calculated using GraphPad Prism 8.

#### Fab preparation

Fab fragments were overexpressed with a His-tag heavy chain expression vector and co-transfected with a light chain vector in Expi293F cells. Fab fragments were also produced by papain digestion with a Fab preparation kit (ThermoFisher, Cat. 44985) according to manufacturer’s protocol. In brief, 0.5-1.0 mg of IgG1 antibodies were mixed with 125 μl papain resin (ThermoFisher, Cat. 20341) for 5 hours in the digestion buffer provided, containing 20 mM cysteine, pH 7.4. Undigested antibody and Fc fragments were removed by incubating digested products with a protein A column overnight at 4 °C, then collecting the Fab-containing flow-through. Fabs were analyzed by 4-20 % Tris-Glycine SDS-PAGE (ThermoFisher, Cat. XP04200BOX).

Statistical analysis Competition clusters were processed in two steps. The mAbs were first grouped based on binding to SARS-CoV-2 subdomains (RBD, NTD, S2). These mAbs, together with ungrouped mAbs, were then clustered based n competition in ELISA or in the cell-based assay, taking reduction of signal by 30% as the competition threshold.

In order to determine the presence of epitope dependent, VH-segment preferential usage, we used resampling to bootstrap p-values. For each cluster with size n, we resampled n VH segments from the observed VH segments in our dataset m times with replacement. P-values were generated by counting the number of resampled clusters for which the frequency of a VH-segment matched or exceeded the frequency observed in the 167 S binders in Data S2, dividing the quantity of these instances by the number of trials m, and performing a Bonferroni correction (e.g. multiplying the p-value by the number of unique VH-segments) (*45*). For these data, the 7 clusters range from 5 to 39 members, and 1 million resampled clusters were generated for each cluster. We also used the same methods to compare the VH-segment composition of each cluster to the VH-segment composition of the general human PBMC repertoire, exchanging the weights of the unique VH-segments with their representation in the averaged general repertoire from 10 healthy controls (*46*).

Public clones were screened from previously reported clones with a total of 616 SARS-CoV-2 related clones (*1, 12, 22, 33*). We also screened 133 Zika (*34*) and 98 Flu (H1N1) (*35*) related clones. Public clones converged on identical VH and VL alleles, with at least 50% identity in CDRL3 and 50% identity in CDRH3.

#### Protein Expression and purification for cryo-EM

Plasmids encoding stabilized variants 2P (*19*) and hexapro (*42*) of SARS-CoV2 S protein were gifts from Jason McLellan (University of Texas, Austin). Spike proteins for electron microscopy were expressed in Expi293F cells grown in Expi293 medium after transfection with spike-encoded plasmid DNA using the Expifectamine 293 transfection kit (ThermoFisher, Waltham, MA). Cells were grown for 6 days before subjecting conditioned media to affinity chromatography following centrifugation and 0.2 μm filtration. The 2P variant of spike was applied first to a Talon cobalt resin (Takara Bio) and eluted with 200 mM imidazole followed by purification over a S200 size exclusion chromatography column (Cytiva). Alternatively, the hexapro variant of spike was applied to a Streptactin resin (IBA Life Sciences), eluted with 2.5 mM desthiobiotin, and used without further purification.

Antibodies were also expressed in Expi293 as described for spike. C12C9 and C12A2 Fabs were prepared by expressing Fabs with a 3C-cleavable histag that was removed using 3C protease (Pierce) following Talon resin purification, as described (*47*). Fabs of C93D9, G32R7, and C81C10 were prepared by papain cleavage of IgG purified by Protein G affinity chromatography and the Fab collected as the flow through from the same resin.

Spike variants and Fabs were exchanged into 10 mM Tris buffer pH 7.5 with 150 mM NaCl for storage at 4°C.

#### Cryo-EM grid preparation

Grids were glow discharged (PELCO easiGlow) for 30 seconds at 15 mA and prepared with a Gatan Cryoplunge 3 by applying 3.5 uL of sample and blotting for 4.0 seconds in the chamber maintained at a humidity between 86% and 90%. Protein complexes were formed with spike and a 3-fold excess of Fab one hour before freezing and applied with without further purification.

Preliminary studies on 2P S protein complexes with C12C11 were performed with 1.2 mg/ml total protein and C-flat 1.2-1.3 400 Cu mesh grids (Protochips). Structures of C12A2 and C12C9 bound with 2P S protein were determined using Quantifoil 1.2-1.3 400 mesh Cu grids and 0.5 mg/mL total protein, the former with 0.1% w/v octyl β-D-glucopyranoside to reduce orientation bias. Structures of C93D9, G32R7, and C81C10 with hexapro S protein were prepared with thick C-flat 1.2-1.3 400 Cu mesh grids and 1.2 mg/mL total protein.

#### Cryo-EM image recording

Images for C12C11 complexes were recorded on a Talos Arctica microscope operated at 200 keV with a Gatan K3 direct electron detector. Images for C12A2, C12C9, and G32R7 complexes were recorded on a Titan Krios microscope operated at 300 keV with a Gatan BioQuantum GIF/K3 direct electron detector. Images for C93D9 and C81C10 were recorded on an FEI Technai F20 microscope operated at 200 keV with a Gatan K2 Summit direct electron detector. Automated recording was with Serial EM (*48*) in all cases. Specifications and statistics for images from each of these complexes are in Table S1.

#### Cryo-EM image analysis and 3D reconstruction

Image analysis for all structures was carried out in RELION (*49*). Beam-induced motion correction of micrograph movies was performed with UCSF MotionCor2 (*50*) followed by contrast transfer function estimation with CTFFIND-4.1 (*51*), both as implemented in RELION. Particles were picked from motion corrected micrographs using crYOLO (*52*). A general model was used to pick particles from datasets collected on the Talos Acrtica and Titan Krios; a specific model was trained to pick particles from F20 micrographs.

Extracted particles were downsampled and subjected 2D classification, two rounds for the Titan Krios datasets. Initial models were prepared, and the best of three was used as a reference for 3D classification with C3 symmetry imposed. For the Talos Actica and F20 datasets, all particles from reasonable classes were combined and subjected to 3D autorefinement and sharpening, yielding final reconstructions at 8-11Å resolution. The Titan Krios datasets for C12C9 and G32R7 required additional rounds of 3D classification and 3D autorefinement to converge to final C3-symmetric, full particle reconstructions of 3.0 and 3.6 Å nominal resolutions for C12C9 and G32R7, respectively, but with much lower resolution for the Fab-bound domains. We therefore carried out local refinement as follows. Particle stacks from the final C3-symmetric, full particle maps were symmetry expanded and back projected to create a new C3-expanded reconstruction. Models of the most similar heavy and light chains were extracted from the protein data bank and combined to create initial models for the Fabs of C12C9 (heavy: 5ggu, light: 6ghg) and G32R7 (heavy: 4qf1, light: 7byr). Fab models and the NTD (residues 14-290 of 7c2l) were docked into the reconstructed maps; the RBD from 7bz5 was also docked into the G32R7 map. These docked PDB models were used to prepare initial masks with a sphere radius of 8 Å using NCSMASK and a soft edge of 5 pixels, added with relion_mask_create. The soft mask was then used to make a background-subtracted, subparticle stack as implemented in RELION. Fab-occupied and well-resolved subparticles were identified with 3D classification without alignment. Further rounds of 3D classification with and without alignment were carried out along with 3D autorefinement to obtain final sharpened maps with resolutions of ~4.0 Å for both C12C9 and G32R7 Fabs. Detailed descriptions of the particle processing are in fig. S6 and statistics, including model refinement, are in Table S1. Fourier shell correlations are in fig. S7.

**Fig. S1.**
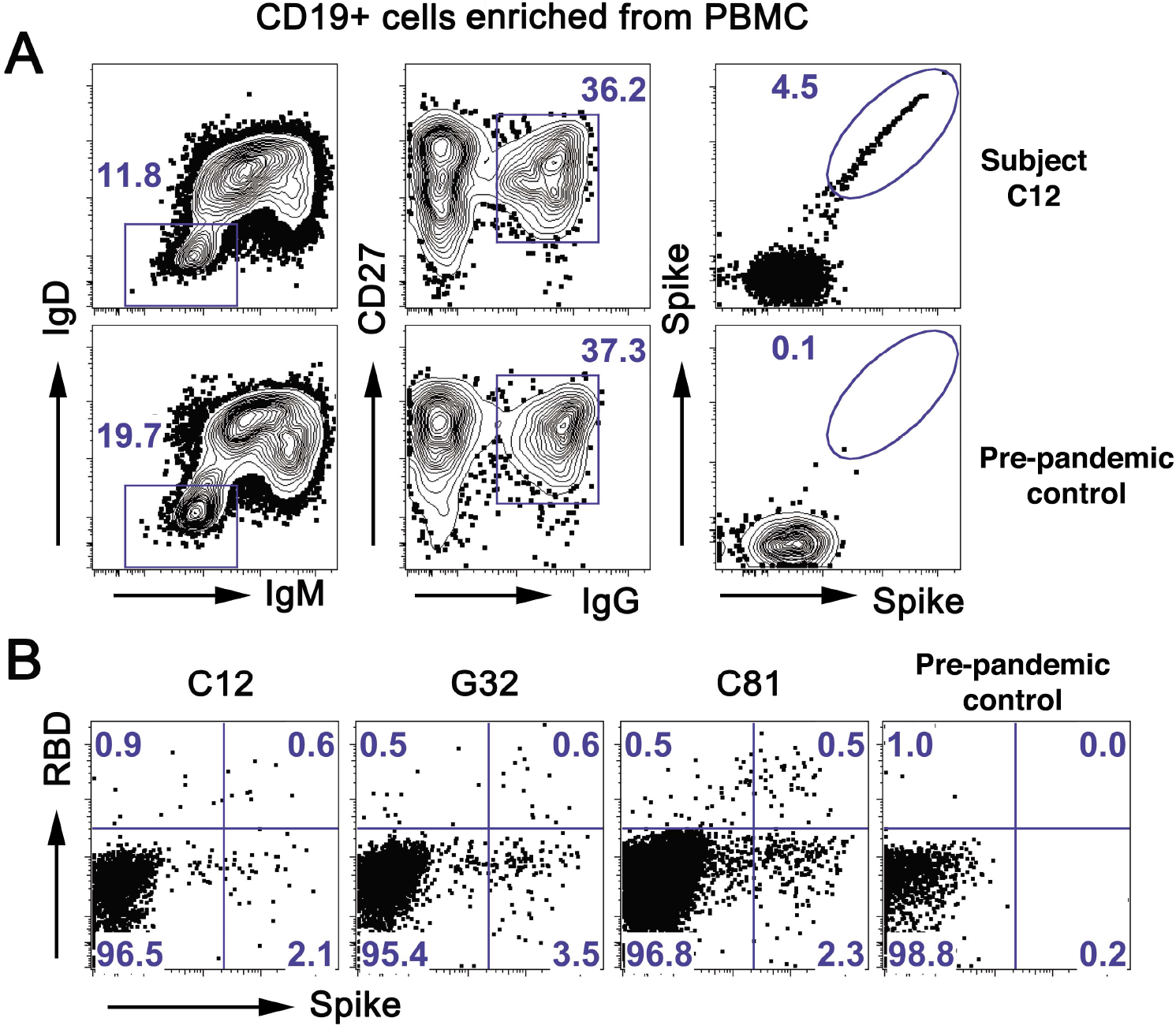
Sorting strategy for SARS-CoV-2 specific memory B cells. (**A**) Representative flow cytometry plots showing CD19^+^, CD27^+^, SARS-CoV-2 spike-binding B cells from a convalescent subject (C12, top row) and a pre-pandemic control (bottom row). PBMCs were pre-enriched with CD19 magnetic beads then gated on live IgD^-^IgM-IgG^+^CD27^+^ and finally on spike (**B**) Representative flow cytometry plots showing spike-positive, RBD-negative B cells for three convalescent subjects and a pre-pandemic control, sorted as in (A) except for the spike gate.

**Fig. S2.**
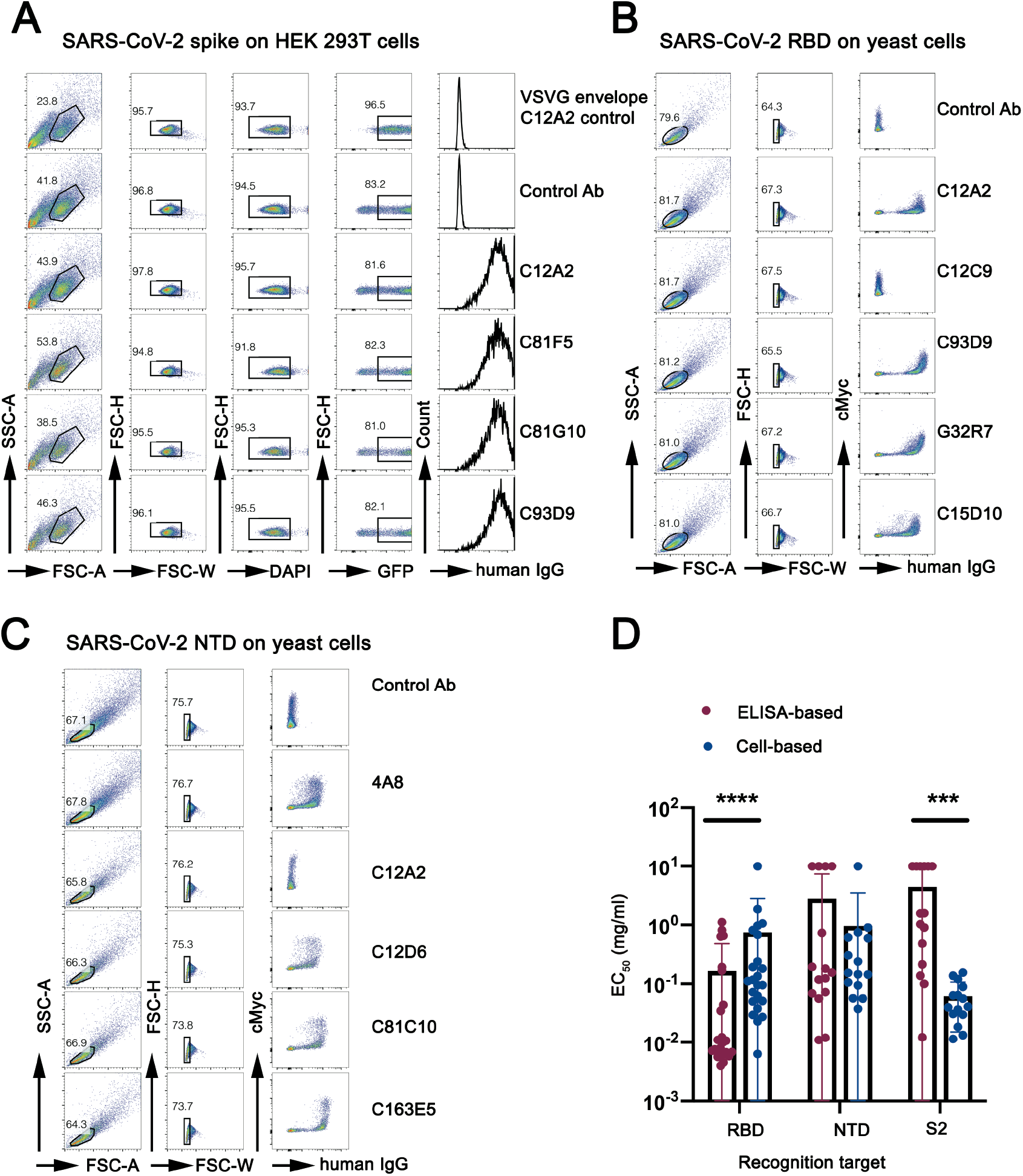
mAb binding to SARS-CoV-2 S, RBD and NTD in cell-surface assay and EC_50_ from ELISA-based and cell-based assay. **(A)** Representative flow plot of mAb supernatant bound to SARS-CoV-2 S on HEK 293T cells. Cells were gated on DAPI-GFP^+^ population. **(B)** Representative flow plot of mAb supernatant bound to SARS-CoV-2 RBD on yeast. cMyc tag indicated yeast that expressed RBD. **(C)** Representative flow plot of mAb supernatant bound to SARS-CoV-2 NTD on yeast. cMyc tag indicated yeast that expressed NTD. See Fig. 1C for the screening color scheme. **(D)** Bar graph of EC_50_ of antibodies targeting RBD, NTD and S2 using ELISA-based and cell-based assay. RBD (n=23), NTD clusters (n=15) and S2 (n=15). *\*\*\*P* < 0.001, \*\*\*\**P* < 0.0001; Paired nonparametric t-test. Data are mean values ± SEM.

**Fig. S3.**
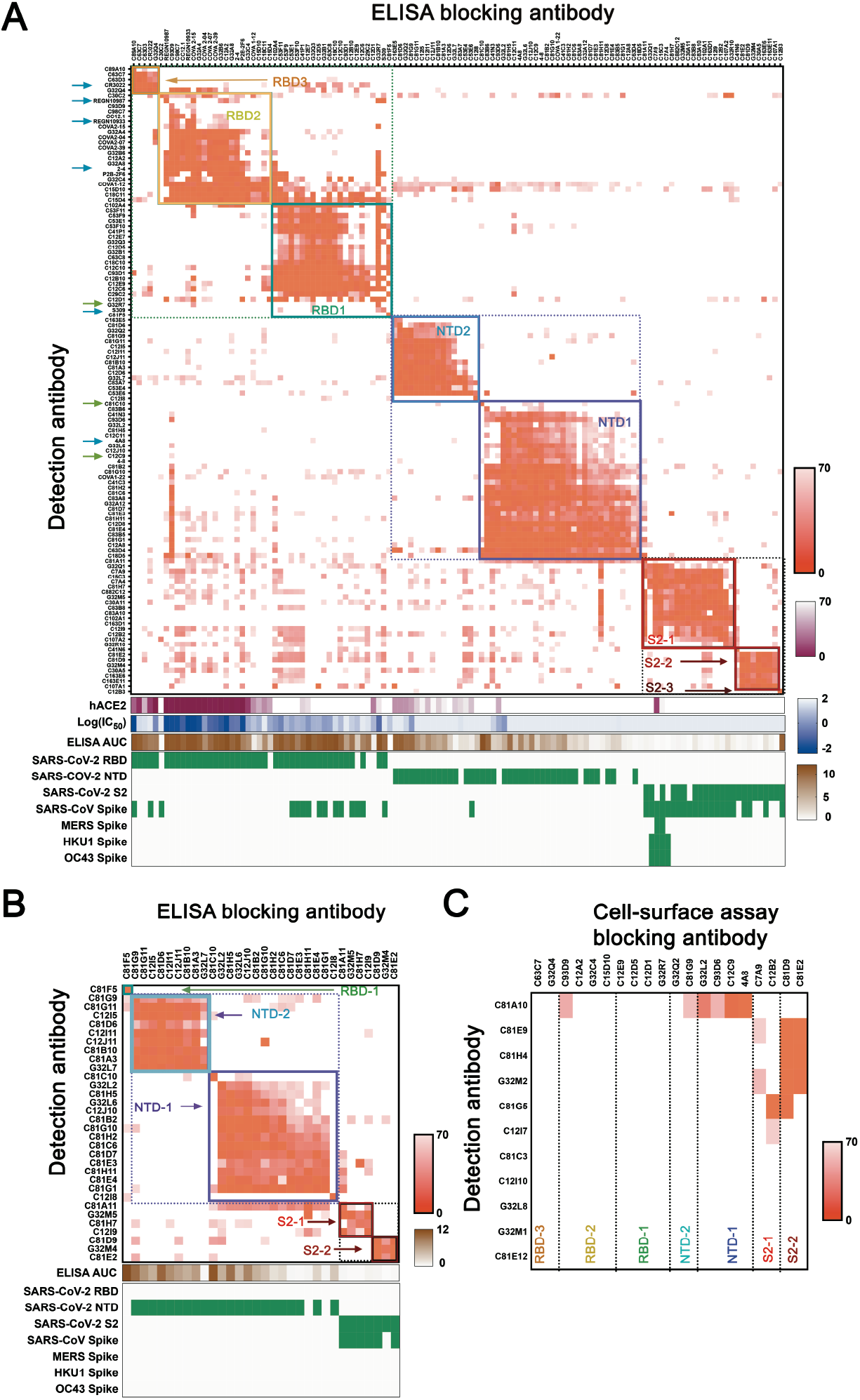
Competition epitope mapping including antibodies from spike^+^RBD-sort and antibodies with published structures. (**A**) Cross competition matrix by ELISA-based competition. Including antibodies from cells gated as spike^+^RBD^-^ increased representation of NTD and S2 clusters. Color and shading scheme, groups defined by hierarchical clustering, and recombinant protein binding as in Fig 2A. Arrows designate antibodies described in the text, including those reported here (green) and those from published work by others (blue). (**B**) Cross competition matrix for mAbs from spike^+^RBD^-^ sort by ELISA-based competition. (**C**) Competition in cell-based assay, for antibodies with binding in ELISA format too weak for reliable blocking measurement. See Fig. 2B for procedures, heat-map color scheme, etc.

**Fig. S4.**
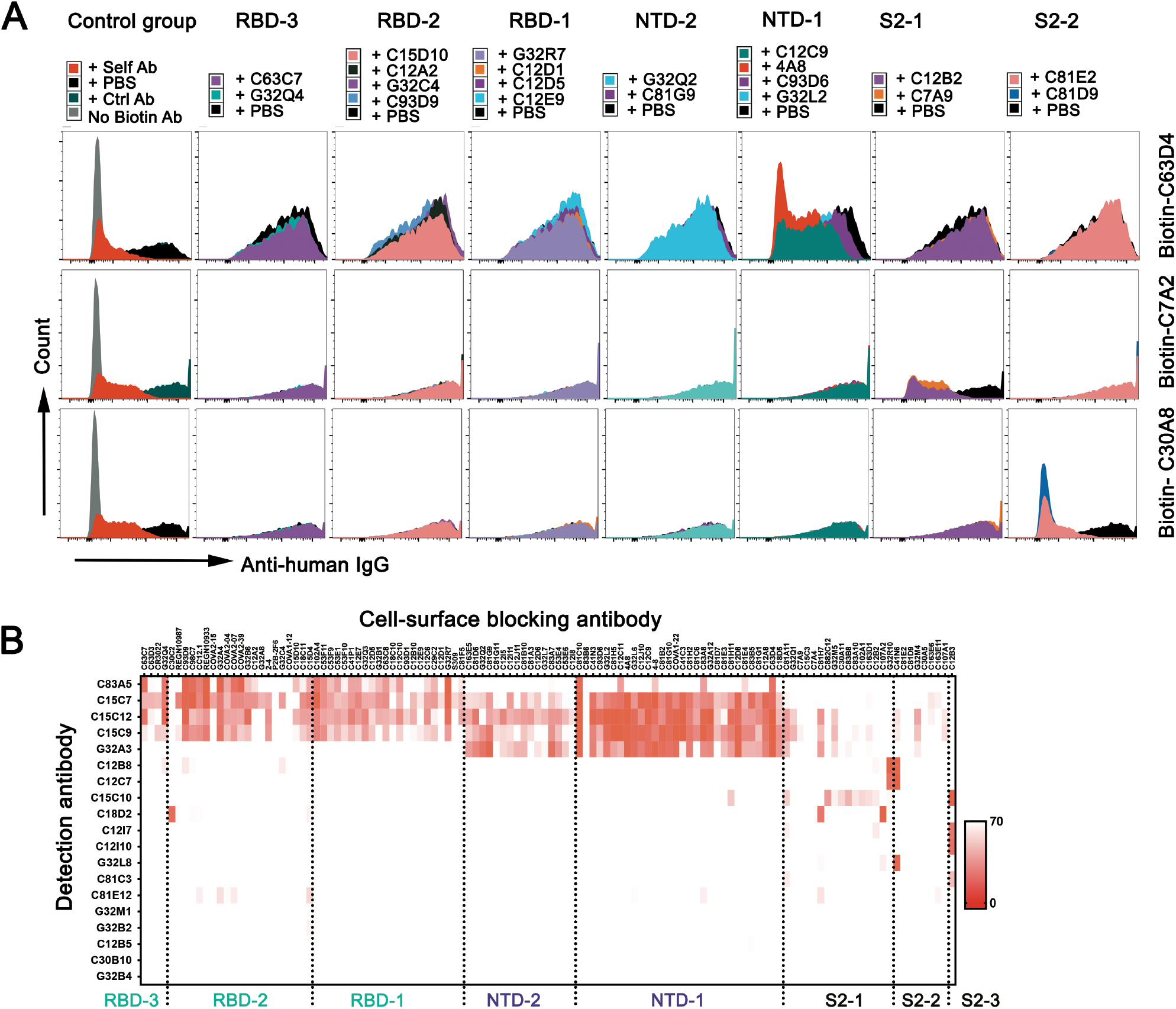
Cell-based competition assay. (A) Representative flows plot for competition, by 20 blocking antibodies representing each of the seven principal clusters (from ELISA: Fig. 2A), for binding cellsurface expressed spike protein by 3 biotinylated antibodies. A non-COVID-19 related antibody and a self-blocking antibody were used as negative and positive controls. (B) Heat map of 19 mAbs with hierarchical clustering from cell-based competition assay with 118 blocking antibodies. See Fig. 2B for procedures, heat-map color scheme, etc.

**Fig. S5.**
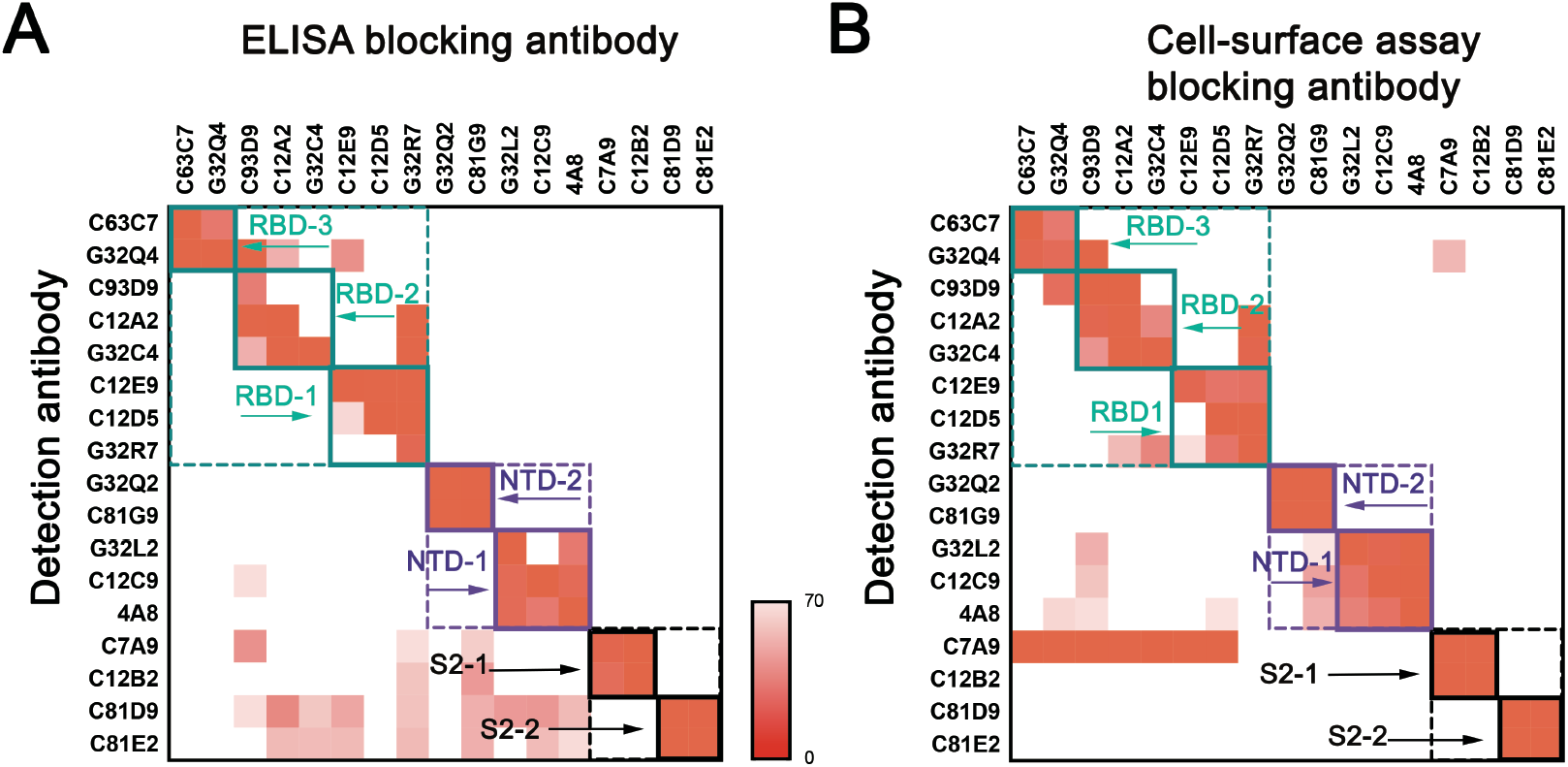
Comparison of ELISA-based and cell-based competition assay. (A) Heat map of 17 mAbs with hierarchical clustering from ELISA-based cross-competition. (B) Heat map of 17 mAbs in (A) with hierarchical clustering from cell-based cross-competition. See Fig. 2B for procedures, heat-map color scheme, etc.

**Fig. S6.**
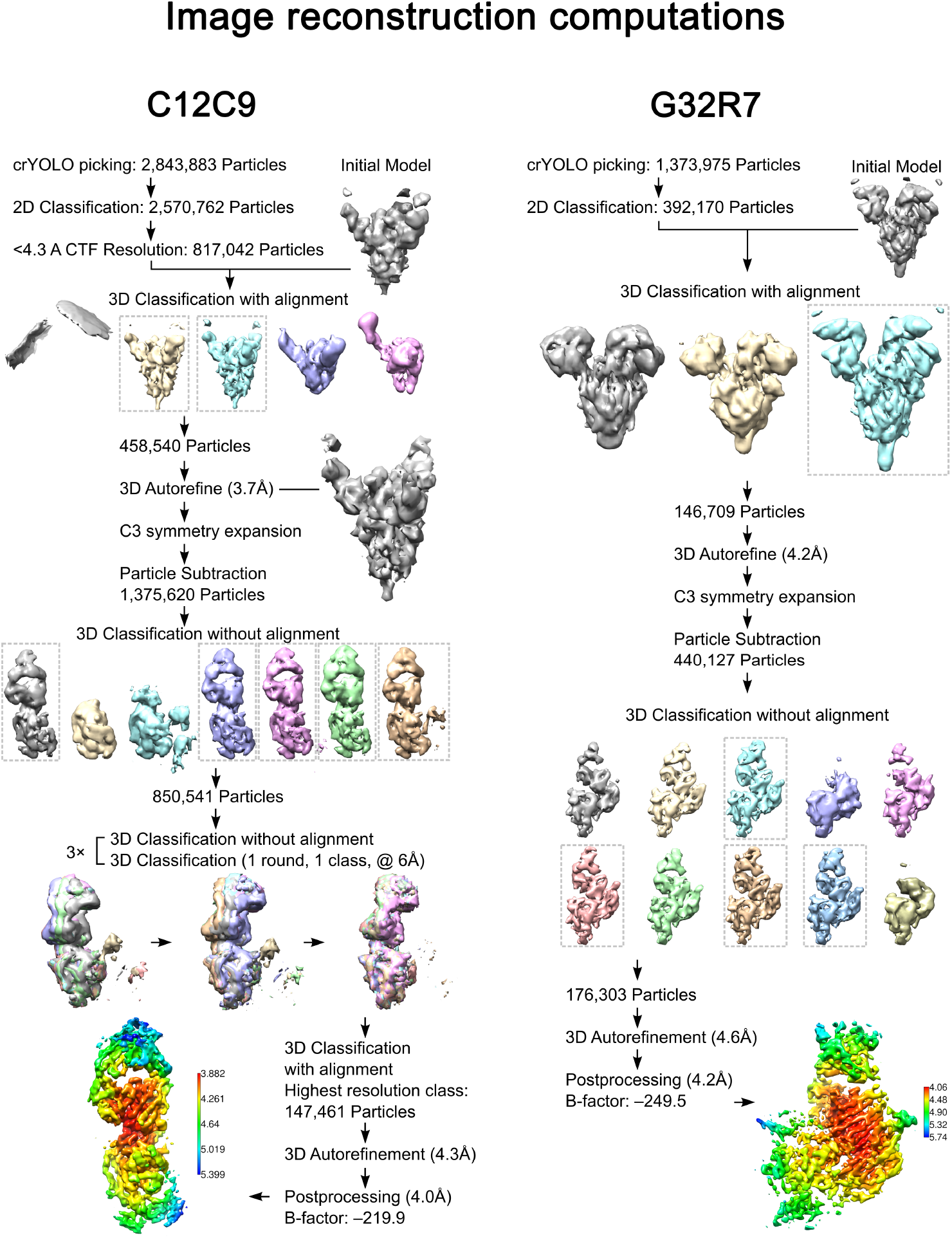
Schemes followed for three-dimensional image reconstructions of C12C9 and G32R7 Fabs bound with SARS-CoV-2 spike ectodomain. See Methods for description of the procedures.

**Fig. S7.**
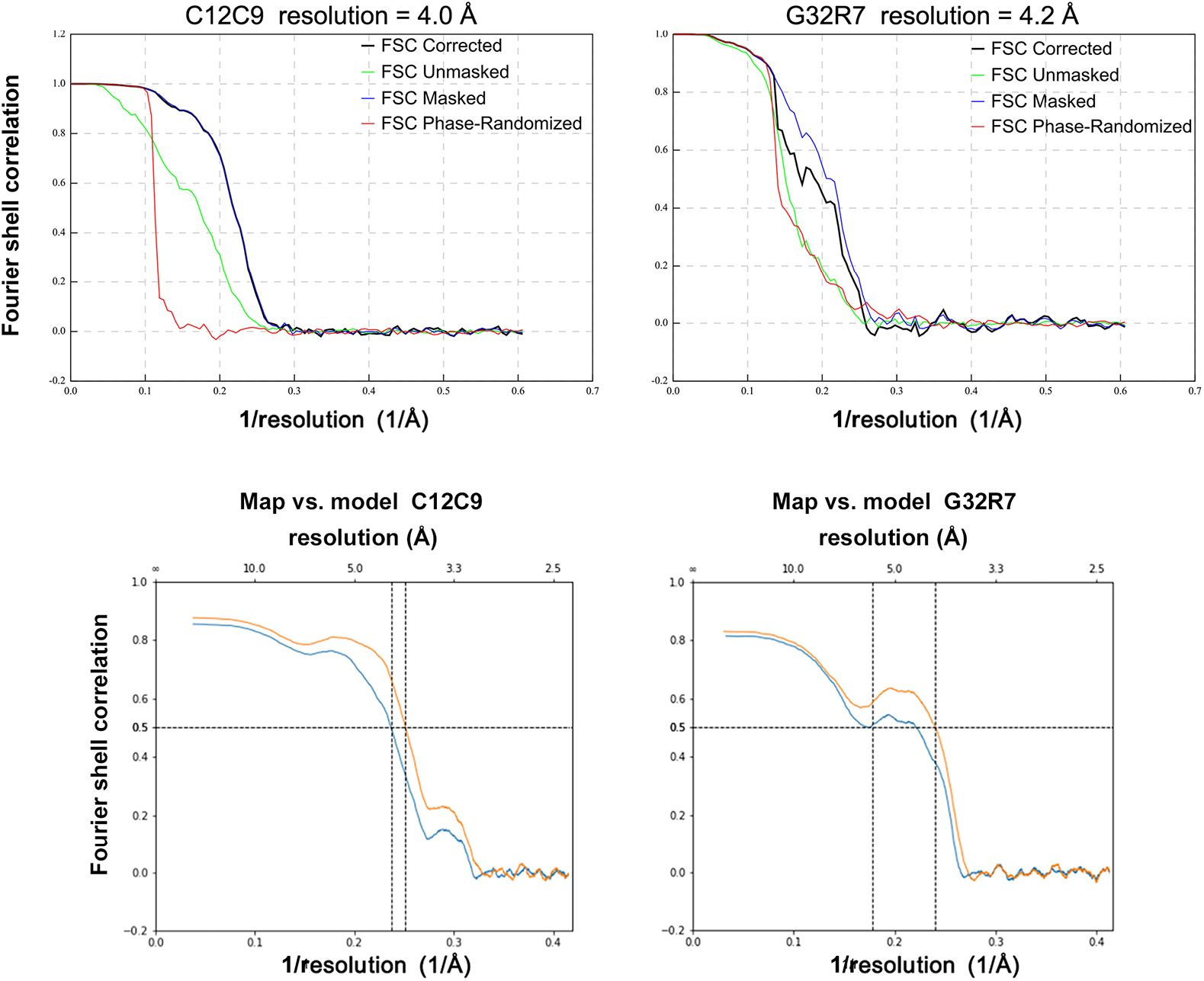
Fourier shell correlation plots for the C12C9 and G32R7 complexes.

**Fig. S8.**
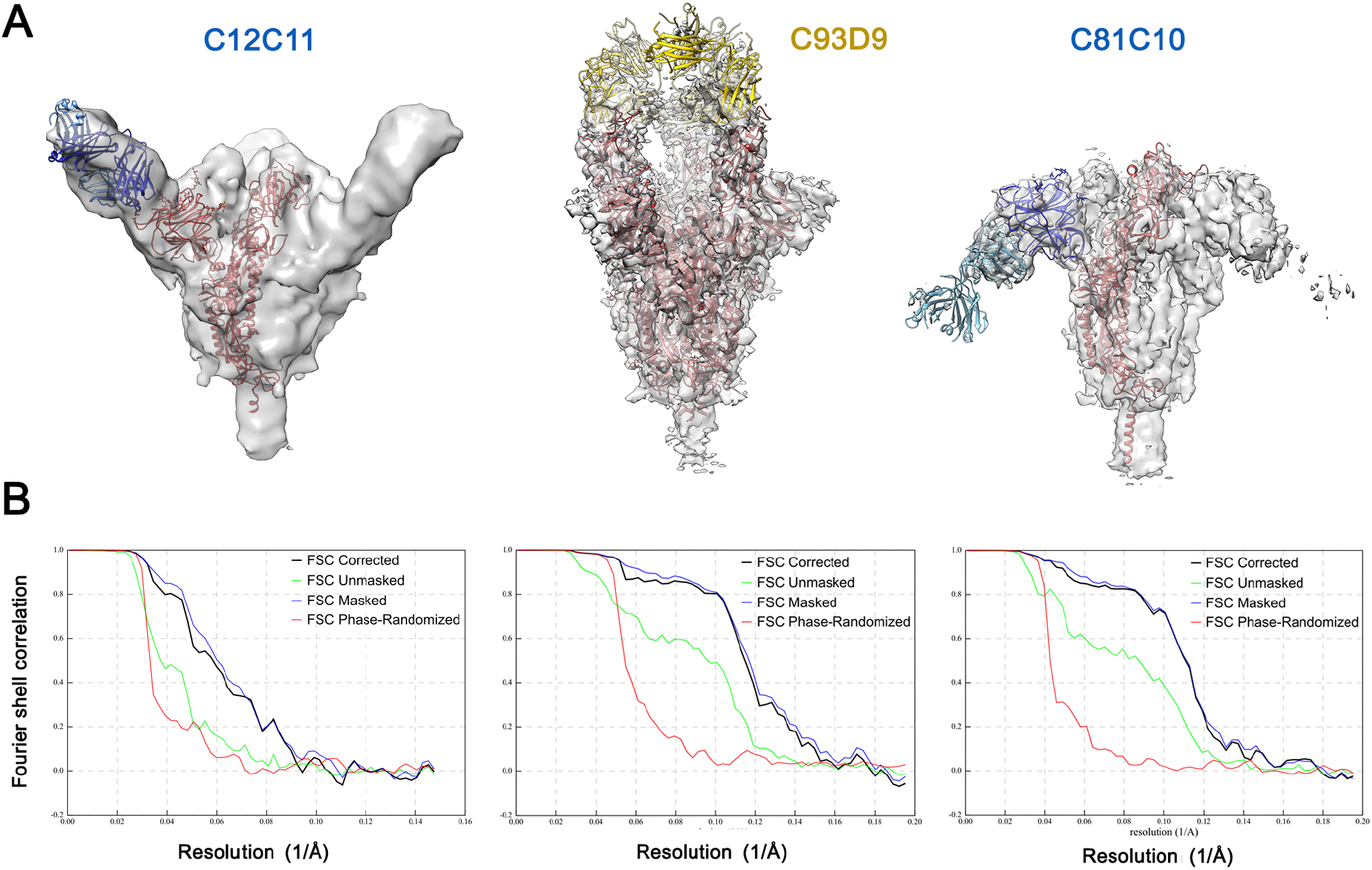
Low- and moderate-resolution structures for C12C11, C93D9 and C81C10 complexes. **(A)** Maps and models. Because the resolution was too low for de novo modeling, we docked the following structures (identified by PDB IDs) into the reconstructed maps, based on similarity of heavy- and light-chain variable domain sequences: 6B0S and 7KJ5 (for Fab and spike, respectively, in C12C11 reconstruction; only a single spike subunit is shown, and the NTD was adjusted manually after 7KJ5 fitting with Chimera “fit-in-map”); 7B3O (for Fab and RBD, docked together, with 6VXX for the rest of the spike, in C93D9 reconstruction); 4QF1 and 6VXX (for Fab and spike in C81C10 reconstruction; only a single spike subunit is shown, and the NTD was adjusted manually after 6VXX fitting with Chimera “fit-in-map”). **(B)** Fourier shell correlation plots for the corresponding maps. The nominal resolutions (0.143 criterion) are 12 Å, 7.1 Å, and 8 Å for C12C11, C93D9 and C81C10, respectively.

**Fig. S9.**
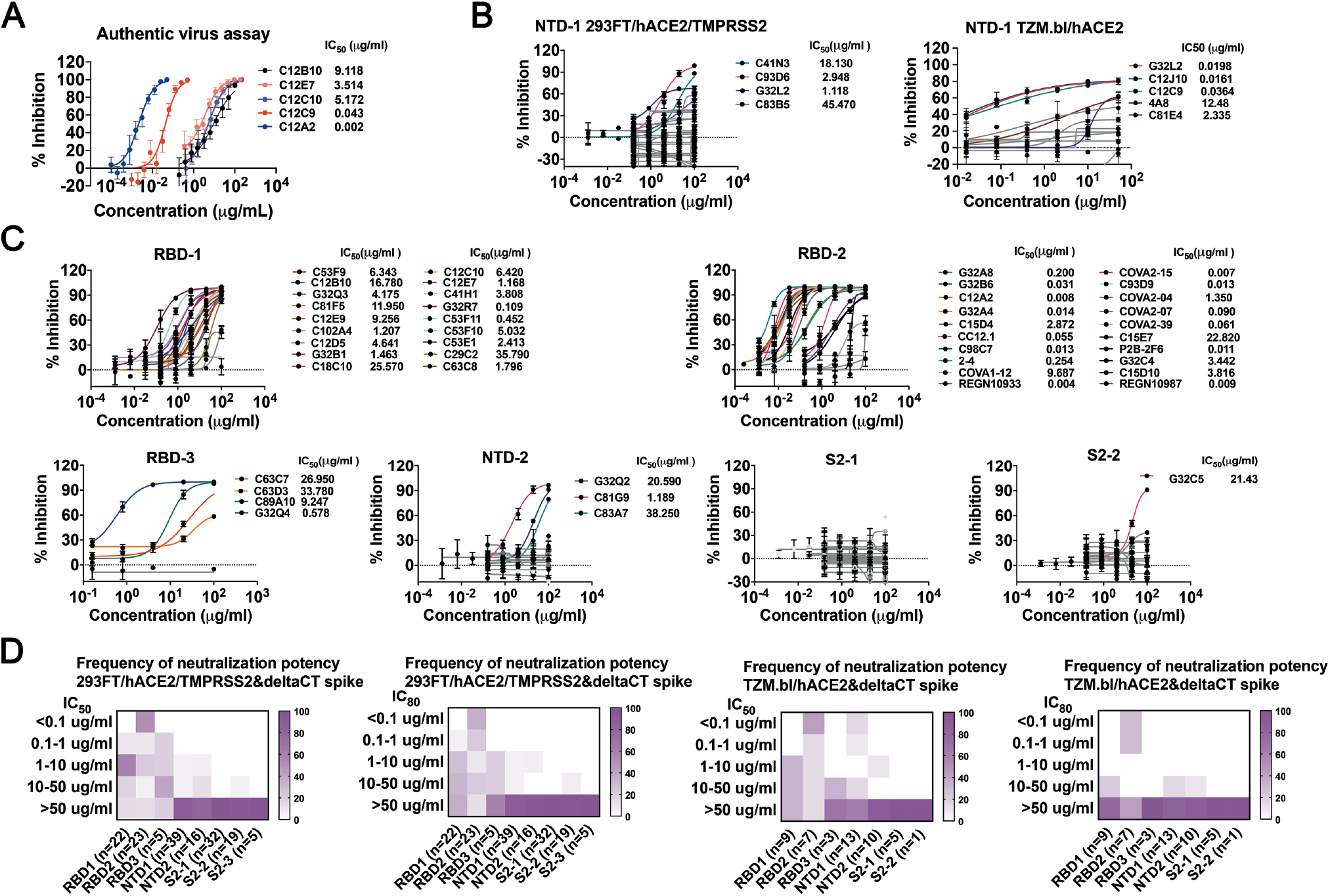
Neutralization profiles for monoclonal antibodies of 7 clusters. **(A)** Authentic virus neutralization profiles of 5 antibodies. **(B)** Pseudovirus neutralization profiles with two cell lines for antibodies from NTD-1 cluster. Neutralization profiles using 293FT cells co-expressing hACE2 and TMPRSS2 were used as target cells shown on the left panel (n=39). Neutralization profiles using TMZ.bl cells expressing hACE2 were used as target cells showed on the right panel (n=13). **(C)** Pseudovirus neutralization profiles for antibodies from RBD-1 (n=22), RBD-2 (n=23), RBD-3 (n=5), NTD-2 (n=16), S2-1 (n=32) and S2-2 (n=19) clusters. Neutralization profiles using 293FT cells coexpressing hACE2 and TMPRSS2 were used as target cells. (D) Distribution of pseudovirus neutralization potency in each competition group. Both IC_50_ (left panel of each pair) and IC_80_ (right panel of each pair) shown, for infection in two different cell lines. Left pair: 293FT cells overexpressing hACE2 and TMPRSS2. Right pair: TZM.bl cells overexpressing hACE2. Color gradient indicates frequency of the clones in each cluster that have the neutralization potency shown by the vertical scale.

**Fig. S10.**
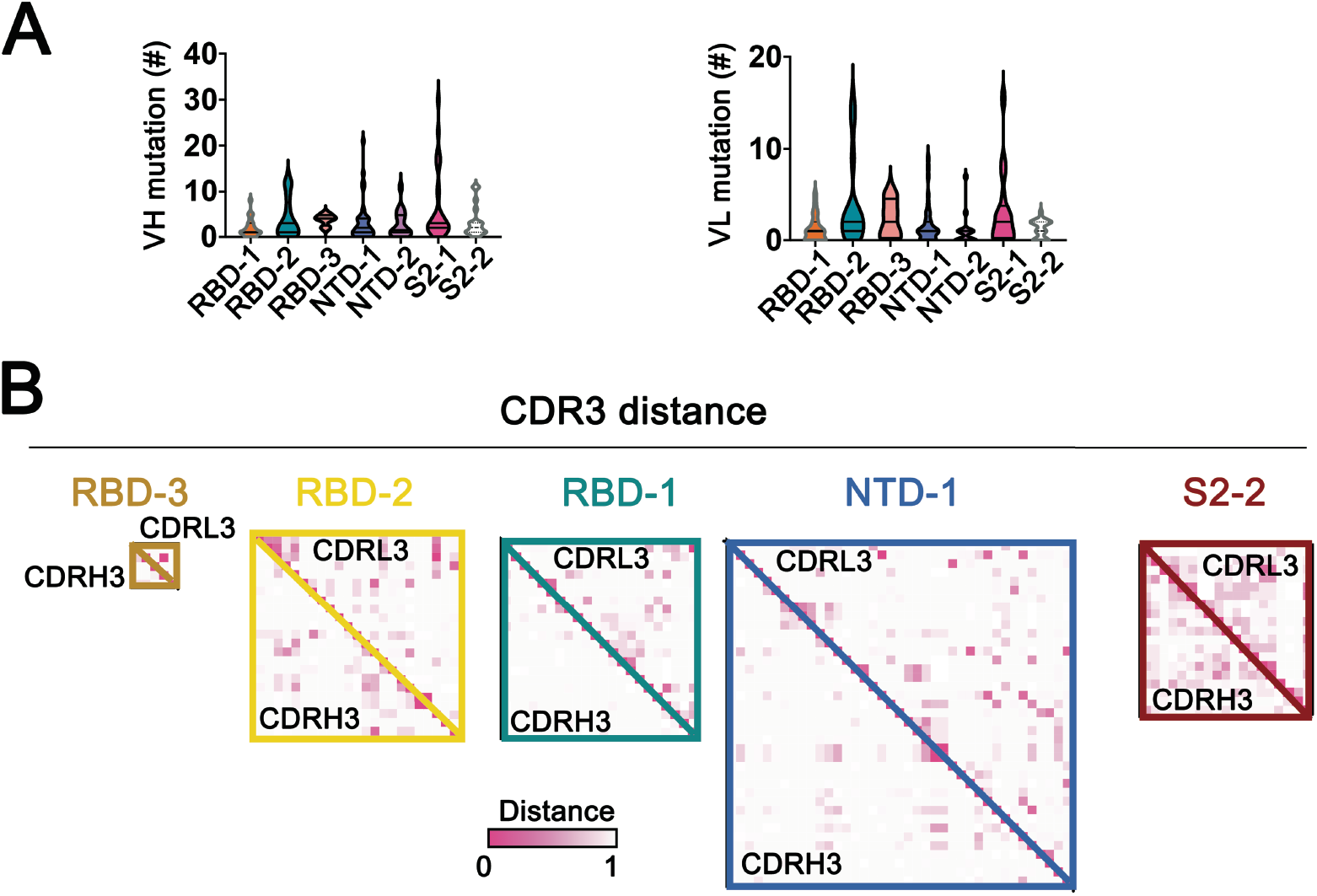
Antibody sequence analyses. **(A)** V(D)J and VJ mutation levels in each of the 7 principal competition groups. Mutations in VH and VL (excluding CDR3) counted by IgBLAST. **(B)** Maps of pairwise distances of CDRH3 (lower left triangle) and CDRL3 (upper right triangle) for the RBD-1, RBD-2, RBD-3, NTD-1 and S2-2 cluster antibodies related to Fig 4B.

**Fig. S11.**
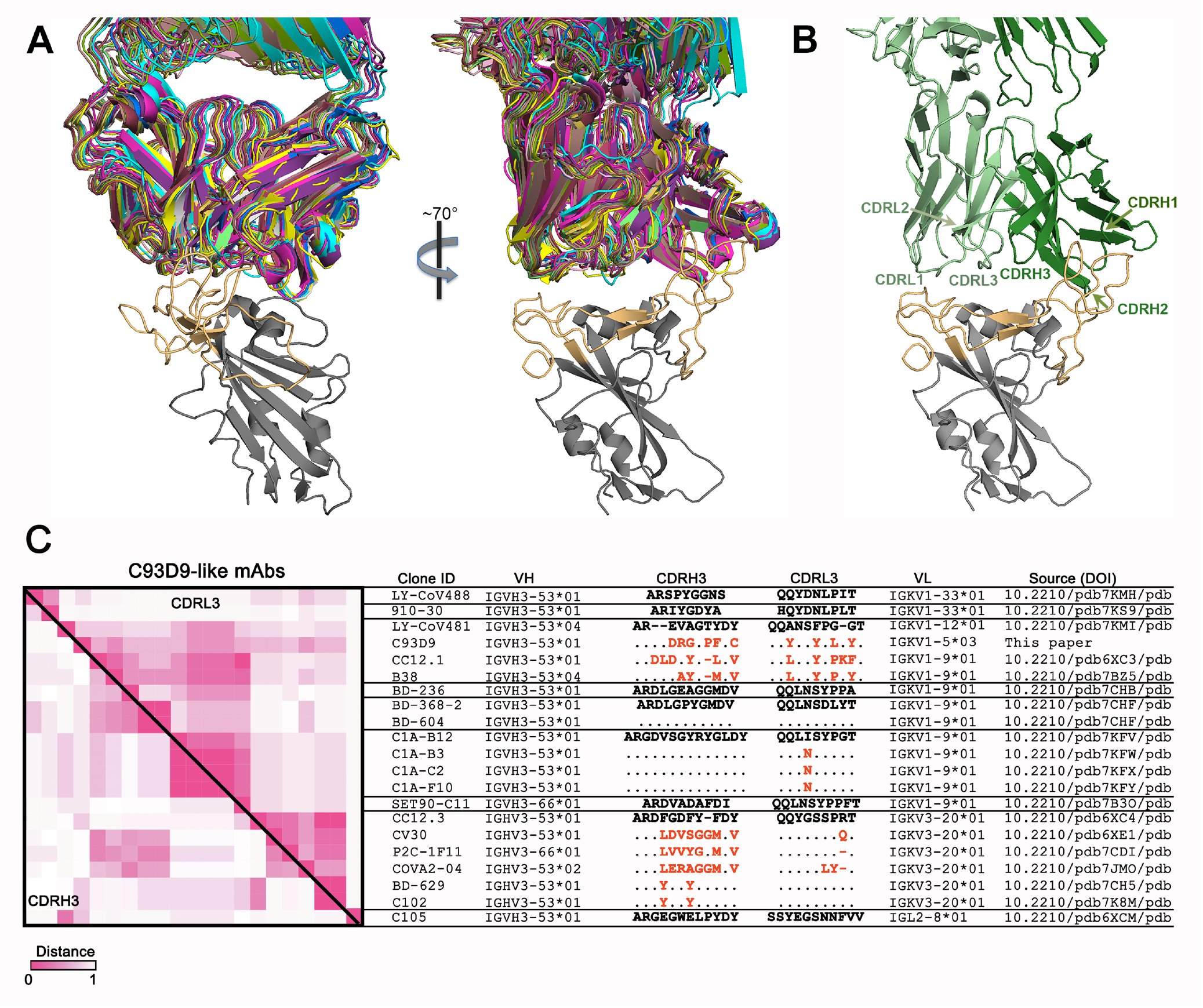
The C93D9 class of antibodies. **(A)** Two views of 20 Fab structures, listed in (C), bound with SARS-CoV-2 RBD. Structures all superposed on the RBD; heavy-and light-chains of each Fab in a distinct color. The figure includes only the RBD from 6YZ5 (not one of the 20), with the RBM in light orange and the rest of the chain in gray. **(B)** View as in the right-hand panel in (A), but showing only the FAB from 7B3O (the closest in sequence to C93D9), with CDRs labeled. The most intimate contacts with RBM residues are from CDRH1, CDRH2 and CDRL1, many with residues constrained in potential variability by ACE2 interaction. **(C)** Maps of pairwise distances of CDRH3 (lower left triangle) and CDRL3 (upper right triangle) for the 21 C93D9 class antibodies in (A) and (B). Pairwise distances analyzed by Mega X. Intensity of color shows the distance, from 0 (identical) to 1 (no identity). The VH and VL genes encoding the antibodies are shown in the indicated groups. Differences in CDR3s from the reference sequences (bold) are in red; dashes indicate missing amino acids; dots represent identical amino acids. IGHV3-66 and IGHV3-53 are very similar VH gene segments, differing by only one encoded amino-acid residue.

**Fig. S12.**
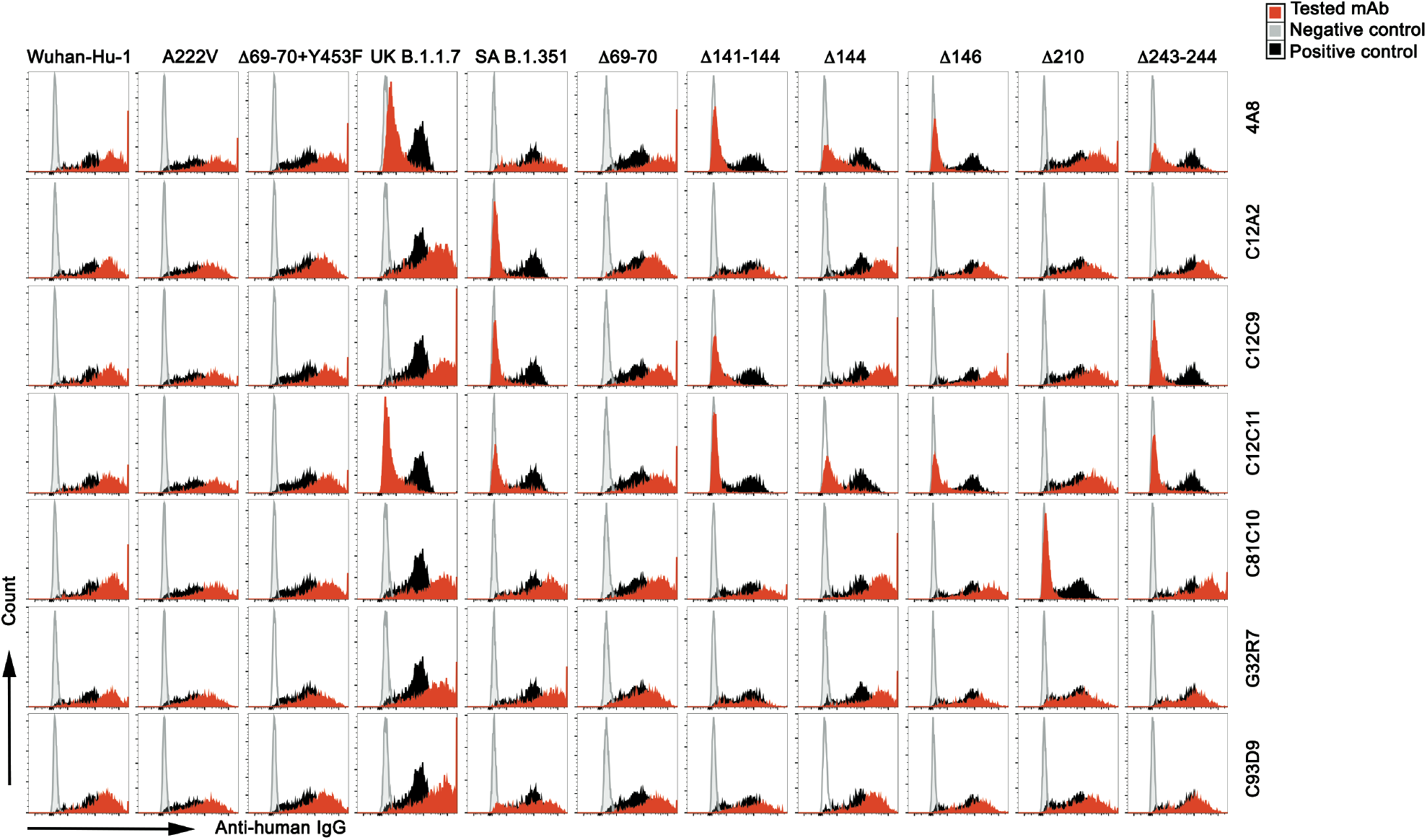
Representative flow plots for mAb binding to the indicated variants. Flow plots for binding of 7 mAbs to Nextstrain cluster 20A.EU1 (A222V), Danish mink variant (Δ69-70 and Y453F), UK B.1.1.7 (Δ69-70, Δ144, N501Y, A570D, P681H, T716I, S982A, D1118H) and SA B.1.351 (L18F, D80A, D215G, Δ241-243, K417N, E484K, N501Y, A701V) and NTD deletion variants. Plasmids with variant S co-expressed with pmaxGFP in HEK 293T cells. Cells were gated on DAPi-GFP^+^. mAb C81E2 was used as positive control, and PBS, as negative control.

**Table S1.** Imaging parameters and model refinement statistics. Electron microscopy imaging statistics and model refinement for spike complexes with Fabs C12C9 and G32R7.

| Table S3. Imaging parameters and model refinement statistics |  |  |
| --- | --- | --- |
| Complex | C12C9 Fab-NTD | G32R7 Fab-RBD-NTD |
| EMDB |  |  |
| Microscope | FEI Titan Krios | FEI Titan Krios |
| Voltage(kV) | 300 | 300 |
| Detector | Gatan K3 | Gatan K3 |
| Magnification (nominal) | 105,000 | 105,000 |
| Energy filter slit width | 20 eV | 20 eV |
| Calibrated pixel size (Å/pix) | 0.825 | 0.825 |
| Exposure rate (e<sup>-</sup>/pix/sec) | 20.784 | 21.296 |
| Frames per exposure | 51 | 51 |
| Total electron exposure (e<sup>-</sup>/Å<sup>2</sup>) | 54.97 | 56.3 |
| Exposure per frame (e<sup>-</sup>/Å<sup>2</sup>) | 1.08 | 1.104 |
| Defocus range (μm) | -1.2, -2.5 | -0.8, -2.5 |
| Automation software | Serial EM | Serial EM |
| # of Micrographs used | 16,812 |  |
| Particles extracted | 2,842,883 | 1,373,975 |
| Total # of refined particles | 147,461 | 176,303 |
| Symmetry imposed | C1 | C1 |
| Estimated accuracy of translations/rotations |  |  |
| Map sharpening B-factor | -219.9 | -249.5 |
| Unmasked Resolution at 0.5/0.143 FSC (Å) | 4.4/4.0 | 6.6/4.7 |
| Masked resolution at 0.5/0.143 FSC (Å) | 4.4/3.9 | 4.7/4.0 |
| Model refinement and validation |  |  |
| Amino acids | 713 | 902 |
| Glycans | 2 | 3 |
| RMSD bond lengths (Å) | 0.009 | 0.008 |
| Angles (°) | 1.010 | 1.238 |
| Mean B-factors |  |  |
| Amino Acids | 135.09 | 60.5 |
| Glycans | 127.94 | 83.7 |
| Ramachandran Favored | 84.11 | 83.71 |
| Allowed | 15.60 | 16.07 |
| Outliers | 0.28 | 0.22 |
| Rotamer Outliers | 1.94 | 0.76 |
| Clash score | 25.63 | 34.43 |
| C-beta outliers | 0 | 0 |
| CaBLAM outliers | 6.89 | 8.66 |
| CC (mask) | 0.74 | 0.66 |
| MolProbity score | 2.79 | 2.70 |

---

**Data S1.**

**Sample source information.**

**Data S2.**

**SARS-CoV-2 mAb clustering, neutralization and sequence features.**

